# Sub-Nanometer Precision using Bayesian Grouping of Localizations

**DOI:** 10.1101/752287

**Authors:** Mohamadreza Fazel, Michael J. Wester, Bernd Rieger, Ralf Jungmann, Keith A. Lidke

## Abstract

Single-molecule localization microscopy super-resolution methods such as DNA-PAINT and (d)STORM can generate multiple observed localizations over the time course of data acquisition from each dye or binding site that are not *a priori* assigned to those specific dyes or binding sites. We describe a Bayesian method of grouping and combining localizations from multiple blinking/binding events that can improve localization precision to better than one nanometer. The known statistical distribution of the number of binding/blinking events per dye/docking strand along with the precision of each localization event are used to estimate the true number and location of emitters in closely-spaced clusters.

## Introduction

Fluorescence super-resolution microscopy methods exploit the independent behavior of fluorescent molecules to circumvent the diffraction limit [1, 2]. Single-molecule localization microscopy (SMLM) methods combine the independent and sparse blinking of emitters with direct inference of emitter locations [3–6]. The emitter positions can be used to reconstruct an image with resolution better than the diffraction limit and in practice this resolution is typically a few tens of nanometers [7]. The resolution is fundamentally limited by the localization precision in the inference step, which itself is limited by the number of photons collected in each blinking cycle [8, 9].

The SMLM methods of (d)STORM [3, 6] and DNA-PAINT [10] can generate multiple blinking/binding events per emitter that are randomly spaced temporally throughout the data collection. These multiple localizations are not *a priori* associated to a particular emitter and are often represented individually in a reconstructed super-resolution image. If these localizations could be correctly assigned and combined, it would result in a higher precision estimate of the positions, scaling as approximately 1/sqrt(*N*) where *N* is the number of repeat blinking/binding events (**Supplementary Note 1)**.

Here, we describe and demonstrate a method for Bayesian Grouping of Localizations (BaGoL) that estimates the number of emitters, assigns the localizations to emitters, and combines the localizations to obtain improved precision. BaGoL uses Reversible Jump Markov Chain Monte Carlo (RJMCMC) [11] similar to our approach to the multiple-emitter fitting problem [12]. The Bayesian formalism allows the use of prior knowledge, such as the blinking/binding probability distribution, and the RJMCMC approach allows exploration of models with a variable number of parameters, which here is the number of true emitters.

While Markov Chain Monte Carlo (MCMC) is confined to problems with a fixed number of parameters, RJMCMC makes inferences about the number of parameters (different models) as well as the parameters themselves. RJMCMC uses an extension to the Metropolis-Hasting algorithm [13, 14] to calculate the transition probabilities of jumping between models. The returned chain can then be used to find the probability distribution of both the models and the probability distributions for the parameters within each model. Our method makes use of RJMCMC to vary the number of emitters in the model and can be used to find the most probable model or to make a weighted average over all models.

The input to the BaGoL algorithm is a set of positions, uncertaintainties and time stamps generated by a traditional SMLM analysis along with a probability distribution for the number of blinking/binding events of a single label. This distribution can be estimated from a dedicated control experiment where the sample is known to be at low labeling fraction [15], from a collection of emitters in the sample that can be identified as single labels, or fiducial markers that have the same characteristics as the sample (e.g., DNA-origami grids). The complete algorithm consists of several steps (**Fig. 1**): 1) Splitting the set of coordinates into smaller subregions to speed up the analysis; 2) Removing outliers; 3) Pre-clustering the data using hierarchical clustering; 4) The RJMCMC algorithm; 5) Generating MAPN and posterior probability results; and 6) Combining the subregions. We briefly describe each step below whereas mathematical details can be found in **Supplementary Note 2**.

**Figure 1:**
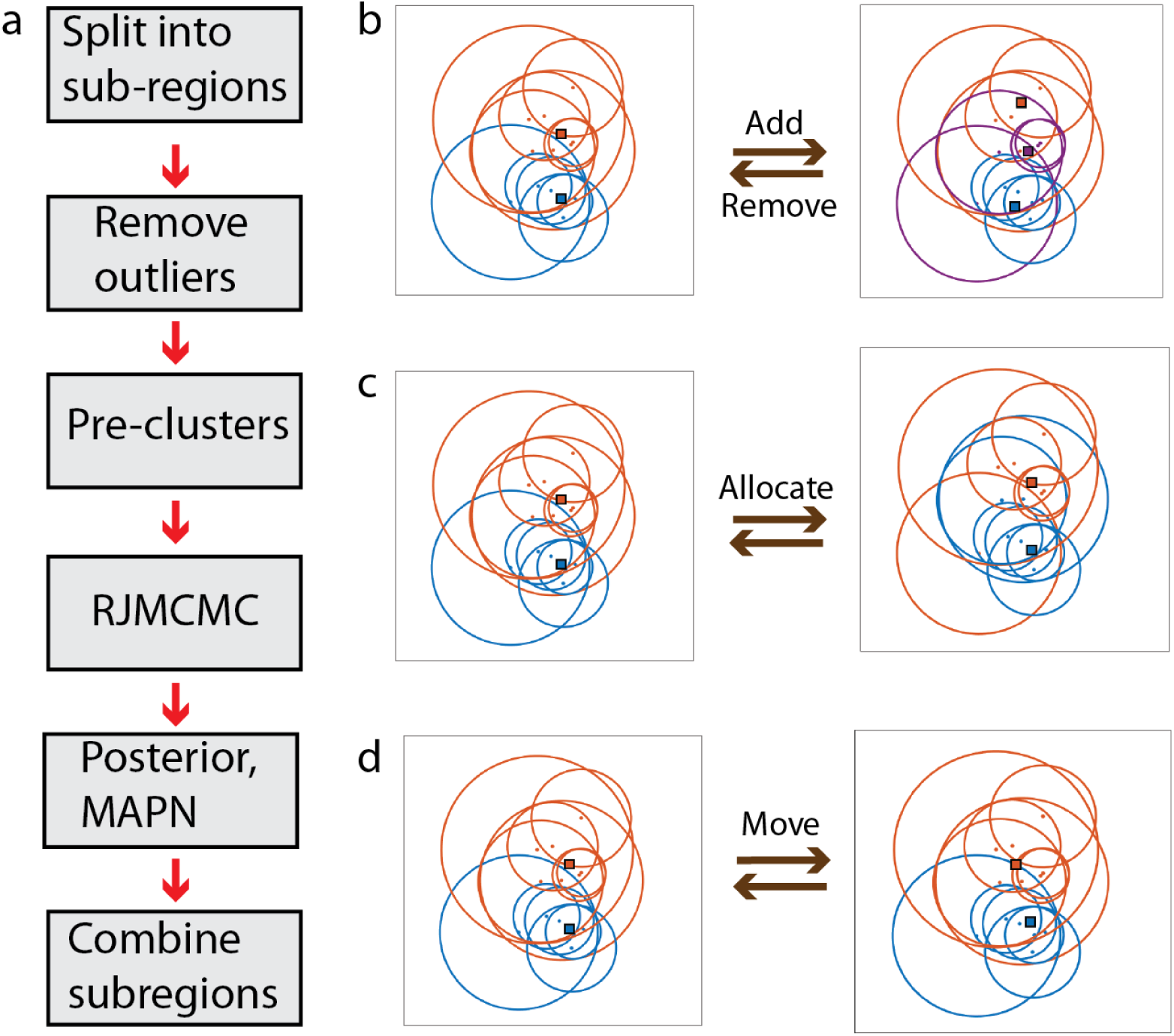
Bayesian Grouping of Localizations Concept and Data Flow. Circles are centered on localizations with radii equal to two times the localization precision. (a) The data flow. The RJMCMC step is illustrated in further detail in (b)-(d). (b) From left to right, a new emitter is proposed in a random location in an ***Add*** jump. From right to left, an existing emitter is picked randomly and is eliminated from the model in a ***Remove*** jump. (c) Localizations are redistributed across the emitters in an ***Allocation*** jump. (d) Given a fixed set of allocations, all emitter positions are updated in a **Move** jump.

1. The data set is split into sub-regions with small overlaps that are used to account for edge effects. The size of the subregions is provided by the user with data density inversely correlated with the subregion size.
2. Localizations that were not generated from a single emitter in the sample are considered outliers and can negatively affect the performance of the algorithm and are therefore removed before analysis. These outliers may arise, for example, from fitting two closely spaced emitters as a single emitter or from non-specific binding of an imaging strand in the DNA-PAINT method. Outliers are identified as localizations with less than *N* other localizations within a distance *R*, or localizations with an intensity higher than two times the mode of the given intensities [16]. *N* and *R* can be chosen from inspecting nearest neighbor distributions.
3. The hierarchical clustering procedure is used to further split the data in the subregions into discrete clusters of localizations that can be analyzed independently. The clusters from the hierarchical algorithm are then sent to the core RJMCMC algorithm.
4. The core RJMCMC step considers the number, *K*, and true locations of emitters along with the assignment of the *N* localizations to *K* underlying emitters treated as a latent variable, *Z*, which is marginalized out in the analysis. We use *θ*={**μ**_1_,**α**_1_ ..,**μ**_*K*_, **α**_K_} as our parameter set, where **μ**_*k*_ is the location of the *k*th emitter and **α**_*k*_ models a linear drift term of the *k*th emitter. The parameter set, *θ*, is explored by proposing and then accepting or rejecting four jump types that are illustrated in **Fig. 1. *Move*** selects new values of *θ* via Gibbs sampling given the current allocations *Z*. ***Allocate*** redistributes all localizations across the *K* emitters. ***Add*** adds a new emitter to the current model and redistributes the localizations. ***Remove*** removes one of the emitters and redistributes the localizations. ***Move*** is always accepted by virtue of Gibbs sampling, whereas the other moves are accepted or rejected with a probability α as given by the rules of RJMCMC. The chain of *θ* is updated, recorded and then used for all subsequent analysis.
5. A posterior probability image of true emitter locations is generated by a histogram-type 2D image of the emitter positions stored in the chain. The model that has the *maximum a posteriori* number of emitters (MAPN) (i.e., the most repeated model) is extracted from the chain and is used to generate an image in the same manner as the posterior probability image. The extracted chain is also used to calculate the MAPN emitter coordinates and their uncertainties using k-means clustering of the distribution of emitter positions in the chain. For both the posterior image and MAPN image and coordinate results, emitter locations that fall into the overlapping regions are removed.
6. The posterior and MAPN images for the subregions are combined to give a single, BaGoL reconstructed image.

## Results

We applied BaGoL to experimental DNA-PAINT SMLM data sets from several structures (**Fig. 2,3**) and realistic synthetic data (**Supplementary Figure 1-11**). **Figures 2a-c** show the results of BaGoL applied to DNA-PAINT data collected from commercially available 20 nm spaced DNA-origami rulers that are intended to be used as test structures. The BaGoL analysis clearly improves upon the traditional SR result and resolves the 20 nm spacing of the ruler with a reported precision of about 1.2 nm (**Supplementary Fig. 12**). The averaging of multiple rulers does not degrade or improve the resolution, indicating that resolution is limited by the data and not sample variability. BaGoL applied to the combined MAPN results over multiple structures (**Supplementary Fig. 12**) gives a ruler separation of 20.7 nm which is consistent with the manufacturer’s specification of 20 nm.

**Figure 2:**
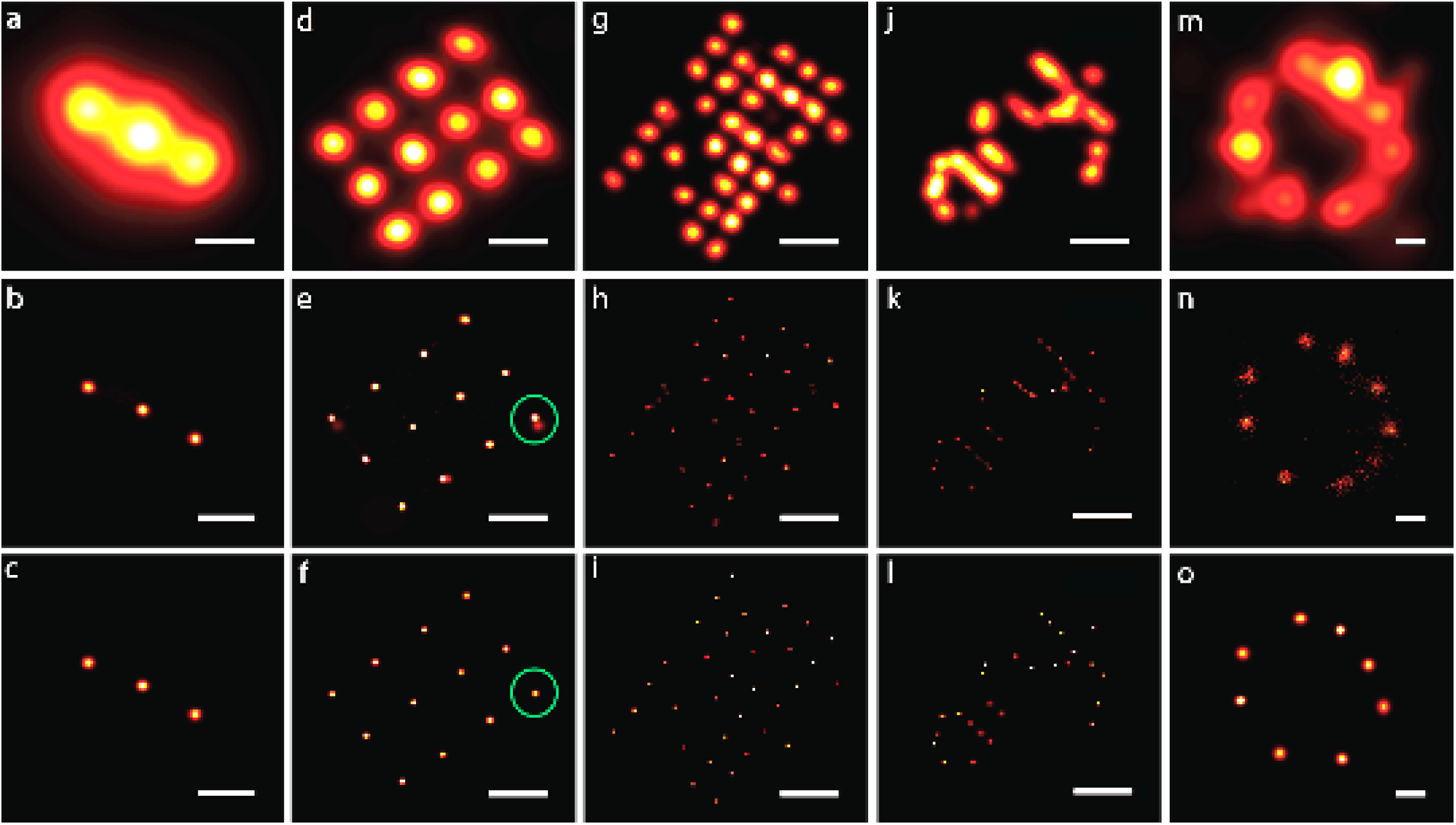
Bayesian Grouping of Localizations applied to various structures imaged with DNA-PAINT. Row 1: Traditional SR analysis with each localization represented by a Gaussian blob of the size of its localization precision. Row 2: Posterior probability image of the chain from BaGoL including all the proposed models. Row 3: The image from the most repeated model (MAPN). (a-c) Gattaquant 20 nm DNA-Rulers; (d-f) Large-spaced DNA-origami grid; (g-i) Small-spaced DNA-origami grid; (j-l) TUD DNA-origami; and (m-o) The Nuclear Pore Complex protein NUP107. The scale bars are 20 nm.

**Figure 3:**
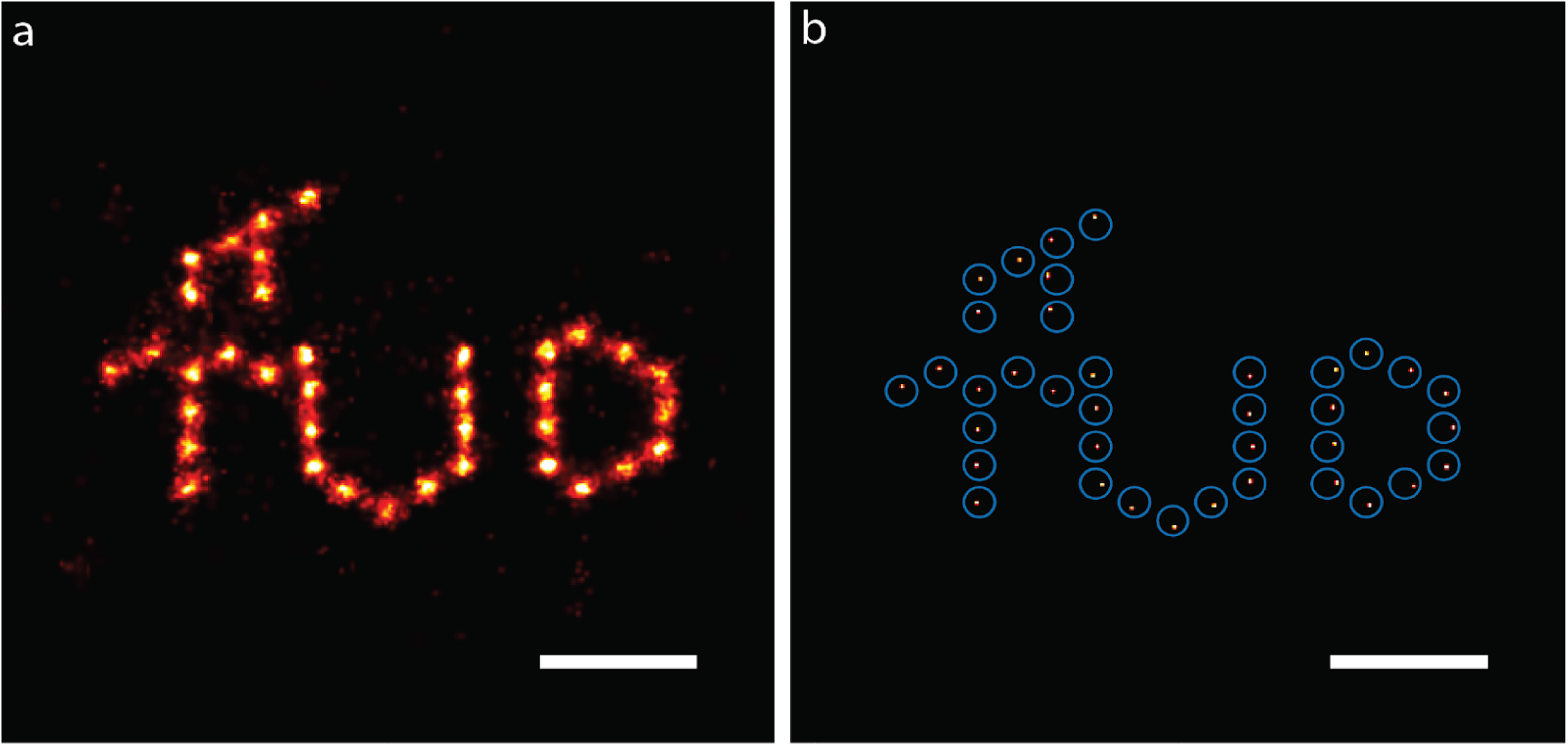
Bayesian Grouping of Localizations applied to aligned structures. (a) The MAPN results of multiple (N=170) structures were aligned with a template and summed. (b) The MAPN image of applying BaGoL to the collection of MAPN results shown in (a). Emitter uncertainties were inflated to match the sample variation and BaGoL. The true emitter locations (template) are shown with blue circles with radii of 2 nm. The scale bars are 20 nm.

**Figures 2d-l** show the analysis of DNA-origami structures that we originally collected and used to demonstrate a template-free particle averaging algorithm for SMLM data [17]. The emitter positions in the ∼ 20 nm spaced array in **Fig. 2d-f** are clearly resolved. The green circle in **Fig. 2e** highlights an area in the posterior image where there was some model uncertainty whereas in the highest probability model (MAPN) image shown in **Fig. 2f** only a single emitter is found. This effect can be seen in several areas of the other structures. In all cases, BaGoL improves upon the traditional SR analysis. **Figures 2m-o** more clearly reveal the 8-fold symmetry of the nuclear pore complex protein NUP107.

For structures that are expected to be similar, BaGoL can be re-applied to aligned outputs from the MAPN coordinates. **Figure 3** illustrates this concept. In **Fig 3a** multiple individual MAPN results were aligned to a template. The emitter positions and uncertainties returned from individual structures were treated like localization positions and uncertainties, and then grouped and combined using BaGoL to generate a high-precision estimate of the average structure (**Fig. 3b**). Note that the sample variation in the TUD structure is larger than the emitter estimation precision. In order for BaGoL to correctly group these, we inflated the uncertainties to match the sample variation. The resulting TUD structure matches that expected from the DNA-origami design [17].

To assess BaGoL’s performance with closely-spaced emitters, we simulated two emitters with separations from 1 to 10 nm (**Supplementary Fig. 1**). In all cases, BaGoL gives an improved representation of the true emitter positions than can be seen in the traditional SR reconstruction. At 1 nm separation, the correct number of emitters is estimated but the position uncertainty is the same scale as the separation and the two emitters are not resolved. At 2 nm separation, the two emitters are resolved with the reported uncertainty matching well with the observed deviation from the true positions. At larger separations, the precision and accuracy improve, which can be explained by less uncertainty in the correct allocations of localizations to emitters.

**Supplementary Fig. 2-6** illustrate the effects of emitter spacing and the average number of localization events on the precision and accuracy of BaGoL for SMLM data of small multimeric structures. **Supplementary Fig. 2,3,4** show the traditional SR reconstruction, the BaGoL posterior image, and the BaGoL MAPN image, respectively. At the smallest radius of 5 nm, BaGoL begins to resolve the ring structure that is not visible in the traditional SR reconstructions, however, the number of emitters in the MAPN image is underestimated in this case. With the larger spacings, BaGoL resolves each emitter with the precision improving with the number of localizations per emitter. The reported precisions from BaGoL and the accuracies calculated by the deviation from the true position improve with number of localizations and spacing and are better than 1 nm for several conditions (**Supplementary Fig. 5,6)**. The convergence of the chain to the stationary distribution is weakly dependent on the data, but has largely become stationary for all cases by ∼1,000 iterations (**Supplementary Fig. 7)**. When including an emitter-dependent linear drift in the likelihood model, the precision and accuracy get worse, in practice leading to an approximate factor of two loss of precision and accuracy (**Supplementary Fig. 8)**.

The dependence of BaGoL performance on the input distribution of localizations per emitter was examined using a 10 nm radius 8-mer synthetic data set (**Supplementary Fig. 9)**. The data was simulated with a Poisson distribution of localizations per emitter with an expected number of localizations of λ = 20. The best results occurred when using the correct distribution. With an incorrect input of λ = 40, the analysis resolves only 7 of the 8 emitters. With an incorrect value of λ = 10, the result shows 8 emitters, but the positions are broadened as the algorithm tries to fit multiple emitters to the localizations from each single emitter. Using a broad gamma distribution performs nearly as well as the correct distribution.

We compared BaGoL to several other clustering algorithms using the synthetic 8-mer with 10 nm radius. The results are shown in **Supplementary Fig. 10.** BaGoL identifies 8 emitters with a precision of a few nanometers. The DBSCAN method is sensitive to its two parameters, the maximum separation distance and the minimum number of elements in a cluster. Even when using *a-priori* reasonable values for these parameters, DBSCAN failed to distinguish separate clusters and grouped all localizations into a single cluster. We applied a Bayesian clustering algorithm developed for super-resolution data [19] to the data and this algorithm also did not separate the localizations into more than one cluster (we note that this was not the original intent of the algorithm). The k-means algorithm was given the correct number of emitters, however, the localizations were not partitioned correctly with some emitter locations placed beyond the view of the image. Similarly, the Gaussian Mixture Model algorithm was also given the true number of emitters, but the variable localization precision breaks the underlying assumption of Gaussian clusters and the results do not predict the true emitter locations well.

Although we envision that BaGoL will be most powerful when applied to small clusters of emitters using DNA-PAINT with many localizations per emitter, it can be applied to any SMLM data set. We show results of BaGoL when applied to SMLM data from a very common sample type often used as a test structure for super-resolution imaging - a dSTORM data set of microtubules. We estimated a localizations/emitter distribution from the data by running BaGoL with a wide prior on number of emitters, then using the distribution of localizations/emitter returned from the analysis (mean=3.5) to generate a new prior and reinitiated the BaGoL analysis. The results are shown in **Fig. 4**. The improved precision reveals that the sample is somewhat under-labeled. The parallel tracks expected from the 2D projection of a cylinder are more clearly visible in many areas in the BaGoL result. We also simulated a microtubule-like quasi-continuously labeled sample where the average label spacing (0.7 nm) was less than that of the average localization precision (5 nm) (**Supplementary Fig. 11**). The BaGoL posterior image shows clear resolution improvement over the traditional SR reconstruction. The results using a broad distribution on the localizations per emitter are nearly identical to that of using the correct distribution.

**Figure 4:**
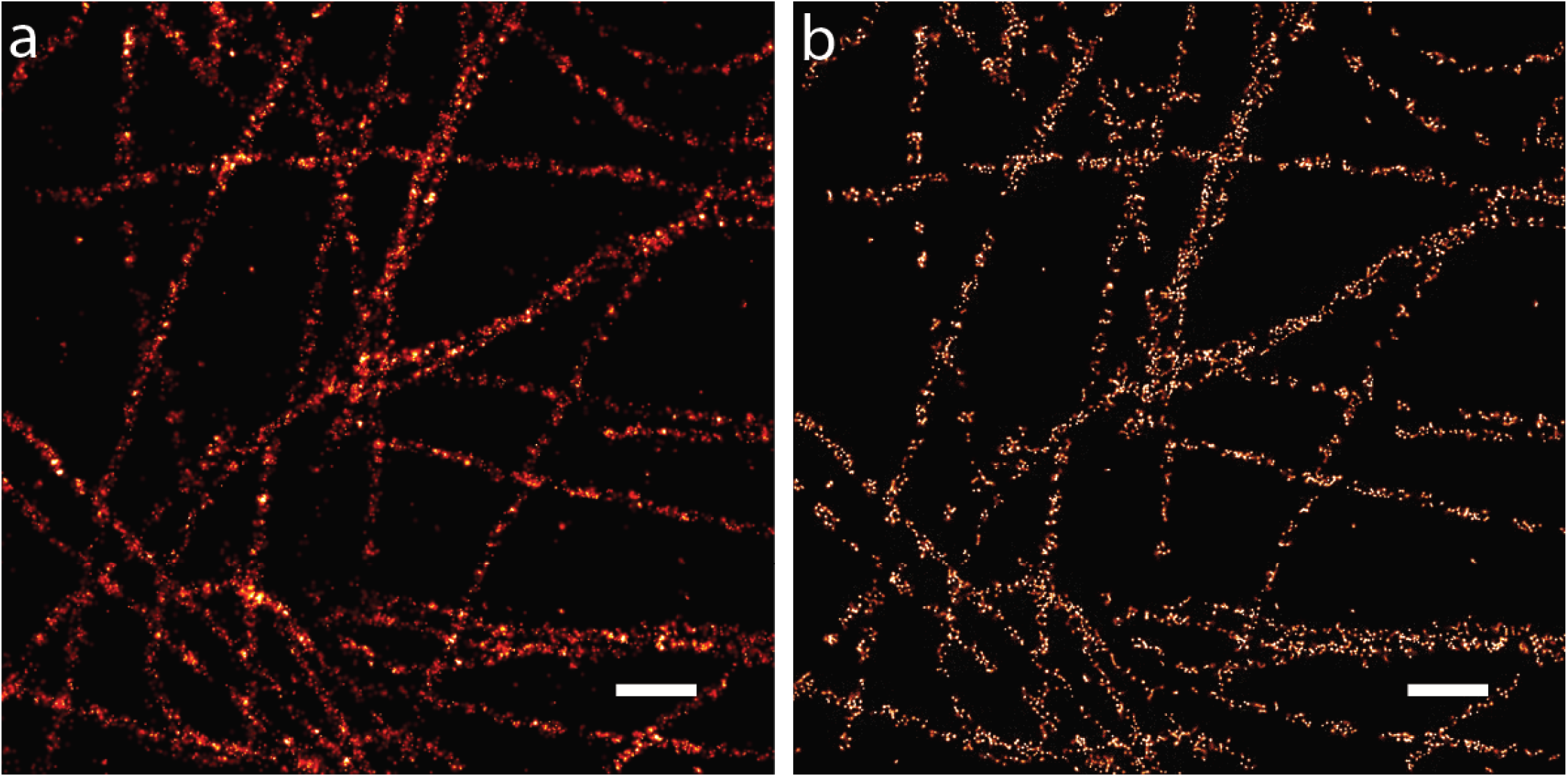
Bayesian Grouping of Localizations applied to dSTORM Data. (a) Traditional SR reconstruction. (b) BaGoL posterior image. Scale bars are 500 nm.

## Discussion

Grouping localizations to improve precision is a simple concept, but requires several things to make it work in practice. There are many clustering algorithms that could be naively employed for this problem such as hierarchical, k-means, Gaussian mixture models [20], DBSCAN [21], Voronoi tessellation [22] and others. None of these general purpose clustering algorithms make best use of the information available in SMLM data, particularly the variable localization precision and prior knowledge of the distribution of localizations from an emitter. We tested several of these algorithms and a naive application of these methods did not produce satisfactory results, as clearly demonstrated in **Supplementary Fig. 10**. Many of these algorithms could potentially be adapted to include this information. We chose the RJMCMC approach described here because the formalism was demonstrated to work well in our past work where we used RJMCMC for the multiple-emitter fitting problem [12] and because the exact number of emitters is not required *a priori*.

The core RJMCMC step of the BaGoL algorithm makes the assumption that localizations are generated by the underlying true emitter, the emitter is labeled by a single dye or docking strand, and that the localization uncertainty is reported correctly. We chose to primarily use DNA-PAINT to demonstrate the quantitative aspect of BaGoL experimentally because it is possible to label proteins with only one docking strand and the binding kinetics are independent of laser intensity (such as might vary across the image or with depth in total internal reflection microscopy) and of buffer conditions such as oxygen or thiol concentration. We also expect that the number of binding events per docking strand will be well described by a Poisson distribution [23]. Experimental SMLM data provides additional challenges because in our experience to date, the SMLM localizations do not always correctly originate from a static, true emitter. Particularly detrimental are 1) high-precision but inaccurate localizations that can arise from fitting two emitters in the raw data as a single emitter; and 2) movement of the emitter during imaging. Preprocessing was essential in removing spurious localizations. Small, nanometer scale movements of individual emitters are clearly present for many emitters in some of the data sets and without modeling this movement, the data is often mis-represented by an excess of emitters. Resolving these issues allowed BaGoL results of experimental data to approach that of synthetic data.

In principle, imaging longer would generate more localizations leading to higher precision. In practice, the precision seems to be limited by sample fixation and nanoscale movements. As a rule of thumb, we would recommend targeting about 50 localizations per emitter with an anticipation of ∼ 1 nm precision.

In this work, BaGoL was applied to 2D data. However, the algorithm can be extended in a simple manner to any number of dimensions, most obviously to include the axial direction. This could provide nanometer precision in the axial dimension comparable to that from interference-based measurements [24]. Applications to other dimensions can also be envisioned, such as overlapping spectral data.

The correct grouping of localizations into emitters makes possible downstream analysis such as cluster analysis of the resulting emitter locations. BaGoL is particularly suited for the quantitative analysis of small oligomers, such as dimers, separated by several nanometers.

Finally, the sub-nanometer precision of BaGoL could be combined with other experimental approaches to generate independence and sparsity on the scale of the nanometer precision, such as multi-color imaging or sequential imaging using orthogonal docking strands. The result would be better than nanometer precision using a relatively simple optical setup where precision and resolution are fundamentally limited only by the sample itself.

## Methods

### BaGoL Implementation

BaGoL was implemented as a MATLAB class. The stages of the algorithm were organized as methods and functions of the class. All the methods, with the exception of the frame connection algorithm, were implemented as MATLAB m-files and require the MATLAB Statistics and Machine Learning toolbox. The frame connection algorithm was written in C++ and was compiled into mex-files that could be used within MATLAB. A desktop computer with an i7, 3.64 GHz CPU was used to process both simulated and experimental data. The algorithm took ∼30 minutes to analyze a 40×40 pixel region of dSTORM data of microtubules, **Fig. 4**, and a few seconds for the single 8-mers shown in **Supplementary Fig. 2-6**

### BaGoL Analysis

The probabilities for the jump types were (P_Move_, P_Allocate_, P_Add_, P_Remove_)=(0.3, 0.3, 0.2, 0.2). We used 1000 jumps for the burn-in portion and 2000 jumps for the post-burn-in portion of the chain, except where otherwise mentioned. The mean number of blinking/binding events, λ, per emitter can be determined using various strategies, described in the following method sections. In general, we would suggest imaging a known DNA-origami structure or fiducial markers simultaneously with the sample to estimate the average number of blinking/binding events per emitter. The filtering parameters to eliminate outliers were adjusted independently for each structure.

#### DNA-Origami

DNA-origami structures of the TUD pattern and grid patterns were part of the same data set, so we used large-spaced grids to estimate the value for λ. Multiple isolated large-space grids were selected manually, and λ was approximated as ∼70 by dividing the number of localizations by the number of emitters in those grids. For the TUD logo and grids with small and large spacings, the outlier localizations with less than 10 neighbors within a distance of 1 nm were eliminated before sending the data to BaGoL. The isolated grids and TUD structures were manually picked and processed by BaGoL using a hierarchical threshold of 15 nm. The localization precisions were inflated by 1.5 nm to compensate for what appeared to be an under-reported localization precision. Docking strand movements were not modeled. The used sub-region size was 20 nm. The MAPN coordinates from BaGoL for 170 TUD structures were aligned with a template generated from origami design (**Fig. 3**). The collection of aligned coordinates and their associated uncertainties were again processed by BaGoL using the mean number of binding events λ=160, which is approximately the number of aligned TUD patterns, and the same hierarchical pre-clustering threshold as before. The precisions were again inflated by 1.5 nm, while this time the localizations with less than 15 neighbors within 2 nm were eliminated.

#### DNA-Rulers

150 isolated DNA-rulers were picked manually and analyzed by BaGoL, adjusting the mean number of binding events to λ=50, obtained the same way as explained in the previous section. The pre-clustering threshold was selected as 20 nm, and localizations with less than 15 neighbors in 10 nm were recognized as outliers. The localization precisions were increased by 2.5 nm and no docking strands drifts were permitted. The sub-region size was set to 100 nm. The MAPN coordinates from the DNA rulers were then shifted and rotated to match a template with the known spacing, **Supplementary Fig. 12**. Not all DNA rulers in the test sample were formed correctly. The DNA rulers that did not match the template, defined as structures where the sum of their nearest neighbor distances with the template were more than 6 nm, were removed from the set of aligned DNA rulers. The collection of the MAPN coordinates from the aligned structures were then analyzed by BaGoL with λ=70, hierarchical threshold of 20 nm, and localizations with less than 10 neighbors within 2 nm were eliminated.

#### dSTORM Data of Microtubules

A broad gamma prior was used to cover the range of the possible number of emitters in each pre-cluster. A typical pre-cluster for this data set had ∼20 localizations where the number of possible emitters ranged from 1 to 20. We employed a gamma prior with the parameters *η*=0.4 and *γ*=10, which covered the mentioned range, see **Supplementary Note 2** for a description of the parameters. The returned average for number of blinking events per emitters was λ∼3.5. We fit the returned distribution to a gamma distribution that was employed to implement a more restrictive prior on the number of emitters with parameters *η*=1 and *γ*=3.5. The localizations with less than 2 neighbors within 30 nm were recognized as outliers. We used a pre-clustering threshold of 50 nm, a sub-region size of 200 nm, 5,000 jumps, no scaling of localization precisions, and no emitter drift. The resulting mean number of blinking events per emitter and the average precision of MAPN coordinates for the second run were λ∼3.4 and ∼6 nm, respectively.

#### NUP107

NPC structure was manually selected, and then processed employing BaGoL. The parameters used for BaGoL were λ=22, obtained using a broad gamma prior as described above, 50 nm for the hierarchical pre-clustering threshold, localizations with less than 4 neighbors in 12 nm were removed, raw precisions were increased by 5 nm, and no movements were allowed for docking strands. The sub-region size used was 400 nm.

### Structure Alignment

Structure alignment and fusion was performed by minimizing the nearest neighbor distances between a template structure and the experimentally obtained structures. An experimental structure was first moved so that its center of mass matched with the center of mass of the template. It was then rotated and translated to minimize the sum of the nearest neighbor distances between the structure and the template vertices using a Monte Carlo approach. The contributions of localizations to the sum with nearest neighbors further than a cutoff distance or localizations with no nearest neighbors were set to the cutoff distance. The length of the chain was 3000 jumps. For the first and second half of the chain, we, respectively, used rotation and translation jump sizes of 1 and 0.1 radian, 0.5 and 0.05 nm. The second half of the chain was then employed to calculate the rotation and translation that minimize the sum of nearest neighbor distances.

### Generation and Analysis of Synthetic Dimers

Two groups of localizations with λ=50, PSF size of 120 nm, mean intensity of 1700 photons and separations of 1, 2, 5 and 10 nm were simulated, **Supplementary Fig. 1**. The number of localizations per emitter and the average intensities of the localizations were drawn from a Poisson and an exponential distribution, respectively, with the given means. The produced data sets were then processed using a pre-cluster threshold of 10 nm, a sub-region size of 200 nm so that all the localizations fit in a single sub-region, no drift, no scaling of the precisions and no filtering. The lengths of the burn-in and post-burn-in chain were 10,000 each.

### Generation and Analysis of Synthetic 8-mers

Synthetic 8-mers with radii of 5, 10 and 20 nm were generated to mimic positions and uncertainties that would result from a standard SMLM experiment. The number of binding events from each emitter was drawn from a Poisson distribution parameterized by the expected number of events per emitter, λ=10, 20 and 50. The time of the event was drawn from a uniform distribution over the simulated data collection time period. For each binding event, the number of photons collected, *I*, was drawn from an exponential distribution with an average intensity of <*I*>=1800 photons. The localization precision of the blinking/binding event for each dimension was calculated using ***σ***=***σ***_PSF_/sqrt(I). The observed location of the blinking/binding events were generated by drawing a value from *N*(*y*,***σ***^2) where *y* is the true position at the time of the event and *N* is the normal distribution. These data sets were employed to make **Supplementary Fig. 2-7**.

The synthetic data were analyzed using the average number of blinking/binding events similar to what we used to produce those data, a hierarchical parameter of 20 nm, and no filtering, no scaling of the precisions and no emitter movements. The sub-region size was set to 200 nm.

The accuracy data in **Supplementary Fig. 6** are the distances of the found MAPN locations from the true locations. These data are compared to the predicted distribution *f(r) = r/****б***^*2*^ *exp(-(r*^*2*^*/2****б***^*2*^)) (magenta curves), in which the parameter *σ* was taken as the precision mean from the corresponding simulation in **Supplementary Fig. 5**. The distributions were scaled to have the same areas under the curve as the accuracy data over the data ranges displayed.

### Convergence Analysis

The chain convergence and mixing was evaluated by running the BaGoL algorithm two times for 10 identical structures and then producing cumulative histogram images of the locations from the states of the two chains. The image cross correlations of the cumulative images of the locations were computed as a function of the number of jumps for each structure and then averaged over the 10 identical structures. When the calculated cross correlations approached one, it meant that the chain has mixed and converged well, **Supplementary Fig. 7**.

### Comparison of Clustering Algorithms

A synthetic 8-mer was generated as described above with a radius of 10 nm and λ=50. This data set was analyzed using 5 different algorithms, **Supplementary Fig. 10**. The data set was processed with BaGoL using the parameters λ=50, pre-clustering parameter of 20 nm, sub-region size of 100 nm and 10,000 jumps. For the algorithm described in Rubin-Delanchy *et al.* [19], we used the recommended value, 20, for the Dirichlet prior and a gamma distribution with a mean of 4 nm, which was the average of the localization precisions. For DBSCAN, the mean number of data points within a group was set to (λ-2sqrt(λ))/2 and the distance parameter ε was adjusted to be the average of the localization precisions. The best result from 10 different runs of *k*-means with 150 iterations and 8 groups is depicted in **Supplementary Fig. 10**. The Gaussian mixture model algorithm was run with 8 groups.

### Double lines Cross

The simulated cross was comprised of double lines with separations of 6 and 12 nm and lengths of 100 nm, **Supplementary Fig. 11**. 70 emitters were placed at random locations along each line, producing an average density of 0.7 emitters per nm. The emitters were synthesized with average intensity, PSF size and mean number of localizations per emitter of 1800 photons, 120 nm, and λ=20. The generated data set was then processed with two different priors: a Poisson prior on the number of emitters obtained with λ=20, and a gamma prior with *η*=0.07 and *γ*=300, see **Supplementary Note 2** for a description of the parameters.

### Experimental Data Collection

#### DNA Rulers

GATTA-PAINT nanoruler slide sample (HiRes 20R, GattaQuant DNA Technologies) were used as purchased. Imaging was done on an Olympus IX71 inverted wide field fluorescence microscope setup as described previously [25]. Fluorescence excitation of the sample was done using a 642 nm laser diode (HL6366DG, Thorlabs). The laser beam was collimated and passed through a multi-mode fiber (P1-488PM-FC-2, Thorlabs), before being focused on the back focal plane of a 1.45 NA oil objective (UAPON 150XOTIRF, Olympus America Inc.). TIRF excitation of the sample was achieved by translating the laser close to the edge of objective back aperture. Fluorescence emission collected from the nanoruler sample was passed through a quad band dichroic/emission filter set (LF405/488/561/635-A; Semrock, Rochester, NY) and a band pass filter (685/45, Brightline) before being detected using an EM CCD camera (iXon 897, Andor Technologies). A total of 100,000, 256 x 256 pixel frames were collected using a 100 ms exposure time. Data collection on the microscope was controlled by custom-written MATLAB software (github.com/LidkeLab/matlab-instrument-control). The raw super-resolution data of DNA rulers was processed by a single-emitter fitting algorithm [9] and thresholded by p-value and localization uncertainty [26]. Localizations from the same binding events were combined using a frame connection algorithm.

#### DNA-Origami

DNA-PAINT data was collected as described in [17]. The raw super-resolution data for the origami grids and TUD logos were analyzed using the BAMF multiple-emitter fitting algorithm [12, 17]. The resulting localizations were then combined across consecutive frames using a frame connection algorithm. Localizations that were not connected across at least two frames were filtered out.

#### Nuclear Pore Complex

The Nuclear Pore Complex (NPC) protein NUP107 was genetically modified with a HaLo tag and labeled with a DNA-PAINT binding strand [18]. Data was collected in the presence of 5 nM imaging strands. The raw image data was analyzed using PICASSO [10]. Localizations were filtered using an intensity threshold of 10,000 photons and then frame connected.

#### dSTORM Microtubules

HeLa cells were plated on a #1.5 coverslip in growth media and were incubated overnight. The cells were washed once with PBS and then fixed by a two-step fixation: 60 seconds in a solution of 0.6% paraformaldehyde, 0.1% glutaraldehyde, 0.25% Triton X-100 in PBS, followed by 1.5 hours in a solution of 4% paraformaldehyde and 0.2% glutaraldehyde in PBS. The cells were then washed twice with PBS followed by 5 minutes in a solution of 0.1% NaBH4 in PBS. The cells were again washed twice with PBS followed by two 5 minute washes in a solution of 10 mM Tris in PBS. The cells were then washed twice with PBS followed by a 15 minute wash in a solution of 5% BSA and 0.05% Triton X-100 in PBS. The cells were washed once with PBS and then labeled for 1 hour in a solution of 2% BSA, 0.05% Triton X-100, and 2.5 μg/mL of anti-α tubulin-Alexa647 (Mouse mAb) in PBS. The cells were then washed three times for 5 minutes each in a solution of 2% BSA and 0.05% Triton X-100 in PBS. Imaging was performed in a standard dSTORM imaging buffer with an enzymatic oxygen scavenging system and primary thiol: 50 mM Tris, 10 mM NaCl, 10% w/v glucose, 168.8 U/ml glucose oxidase (Sigma #G2133), 1404 U/ml catalase (Sigma #C9322), and 60 mM 2-aminoethanethiol (MEA), pH 8.5. Data was collected as described for the DNA-rulers with 16 ms exposure time and a total of 40,000 frames. Weak 405 nm light was used to accelerate emitters out of the dark state.

## Acknowledgements

This work was supported by NIH grant 1R21EB019589 and the New Mexico Spatiotemporal Modeling Center (NIH P50GM085273). We also acknowledge the UNM Center for Advanced Research Computing, supported in part by the National Science Foundation, for providing high-performance computing resources. In addition, we gratefully acknowledge the use of the University of New Mexico Comprehensive Cancer Center fluorescence microscopy core, as well as the NIH P30CA118100 support for these cores. We thank Sandeep Pallikkuth for collecting the DNA ruler data and David Schodt for preparing the microtubule sample.

## Supplementary Information

**Supplementary Figure 1:**
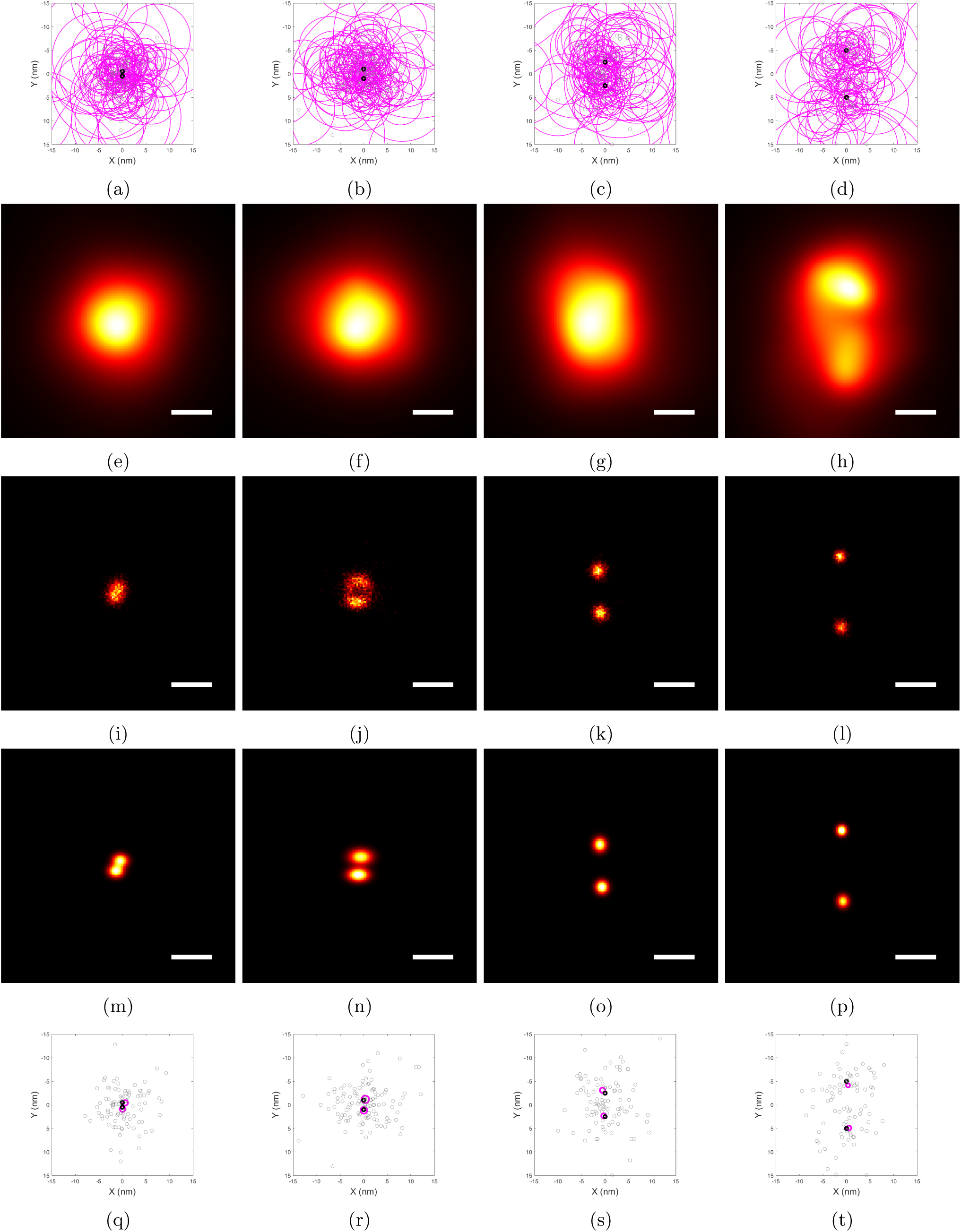
BaGoL applied to two closely spaced emitters. Two emitters are simulated at 1, 2, 5 and 10 nm separations (left to right columns) with an expected *λ* = 50 localizations per emitter and 1700 photons per localization. In all plots true emitter locations are shown as black circles and localizations are shown as gray circles. Magenta circles represent a 1 *σ* localization precision. (a-d) Observed localizations. (e-h) Traditional super-resolution images. (i-l) BaGoL Posterior Images. (m-p) BaGoL MAPN images. (q-t) Plots of MAPN results from BaGoL. Scale bars are 5 nm.

**Supplementary Figure 2:**
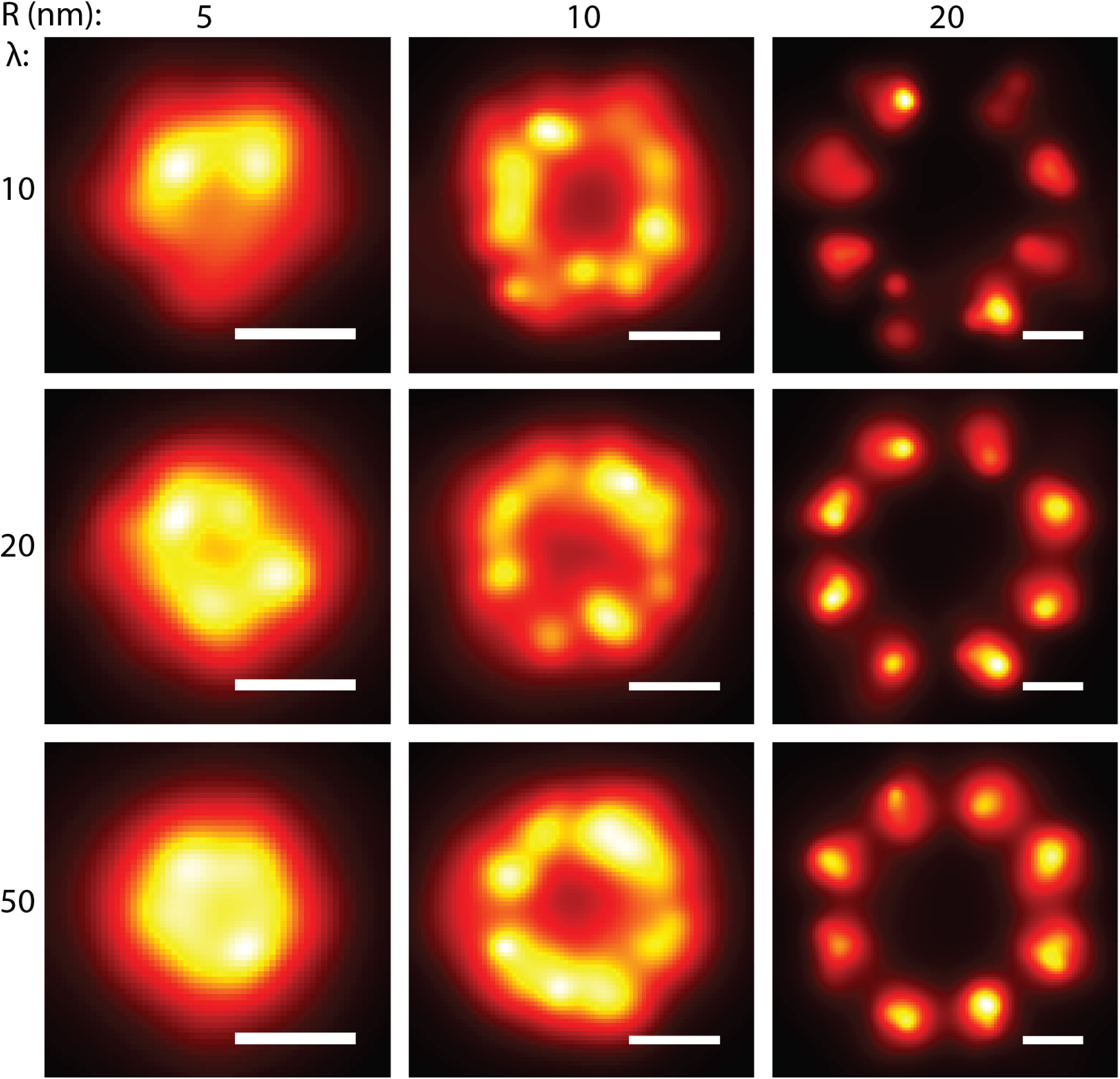
Traditional super-resolution reconstructions of simulated 8-mers. Rows have a fixed average number of blinking/binding events (*λ*) and columns have fixed radii (*R*). The average number of photons per blinking/binding event and the PSF size are, respectively, 1700 photons and 120 nm. Scale bars are 20 nm.

**Supplementary Figure 3:**
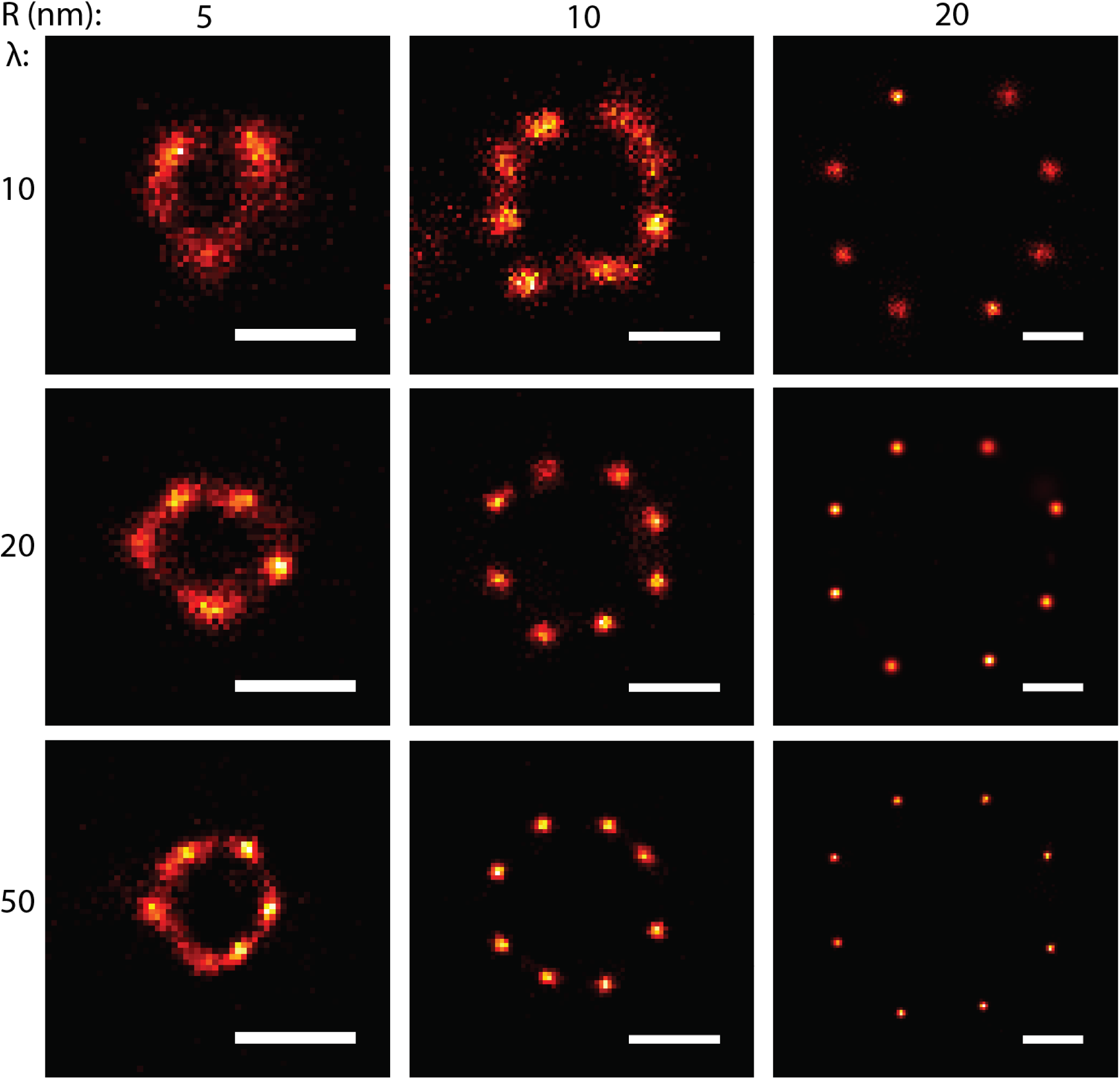
BaGoL posterior images corresponding to the same data shown in Supplementary Fig. 2. Scale bars are 20 nm.

**Supplementary Figure 4:**
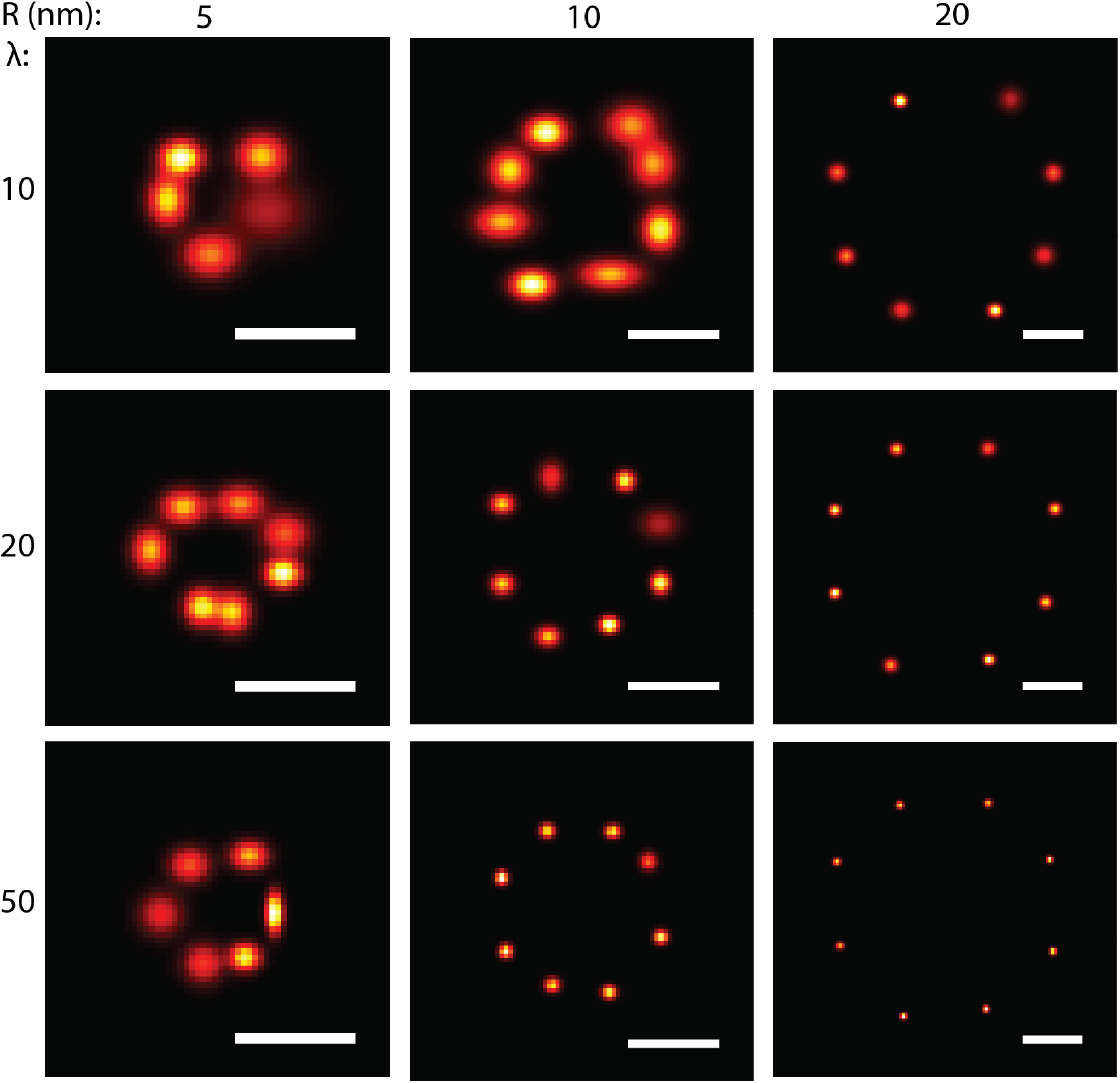
BaGoL MAPN images corresponding to the same data shown in Supplementary Fig. 2. Scale bars are 20 nm.

**Supplementary Figure 5:**
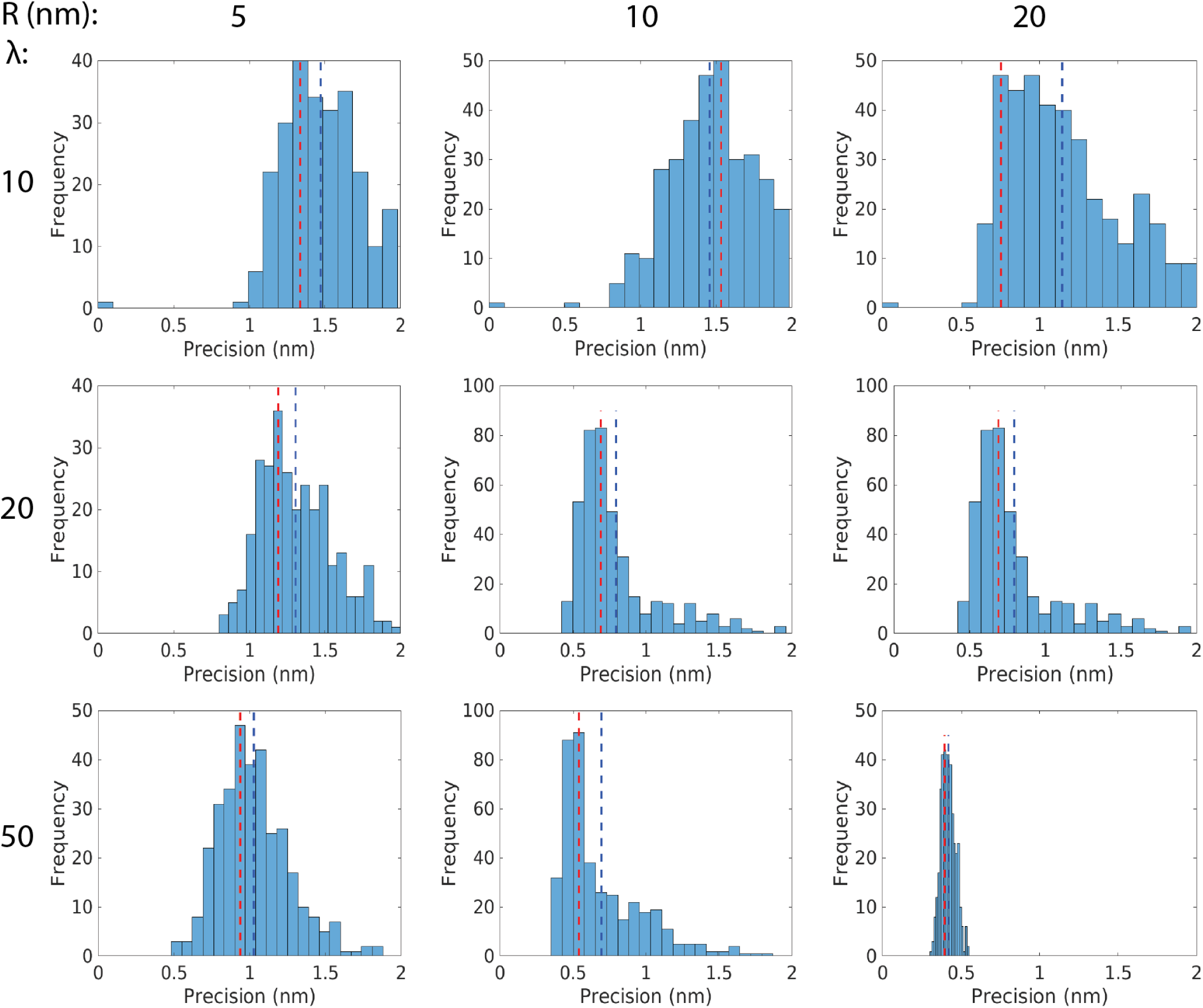
Histogram of precisions corresponding to the MAPN images in Supplementary Fig. 4. The red and blue lines, respectively, show the mode and the mean of the histograms.

**Supplementary Figure 6:**
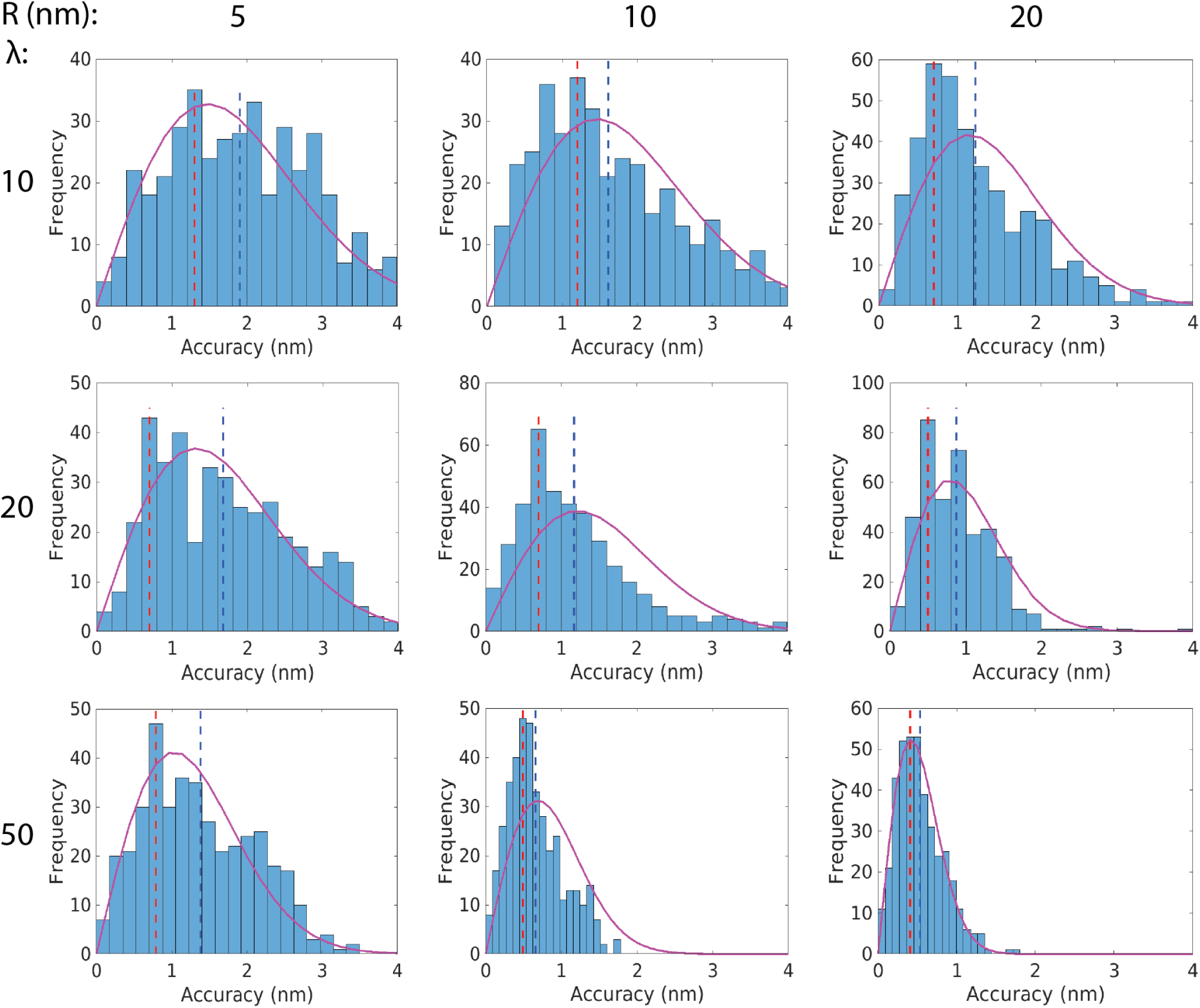
Histogram of accuracies, which is the distance between the real and found locations, corresponding to the MAPN images in Supplementary Fig. 4. The red and blue lines, respectively, show the mode and the mean of the histograms. The magenta curves are fits to the accuracy histograms using *f*(*r*) ∼ *r* exp (−*r*^2^*/*2*σ*^2^), where the parameter *σ* was set to the corresponding precision means, Supplementary Fig. 5. In addition, the areas under the curves for the displayed data ranges were adjusted to match the corresponding histogram areas.

**Supplementary Figure 7:**
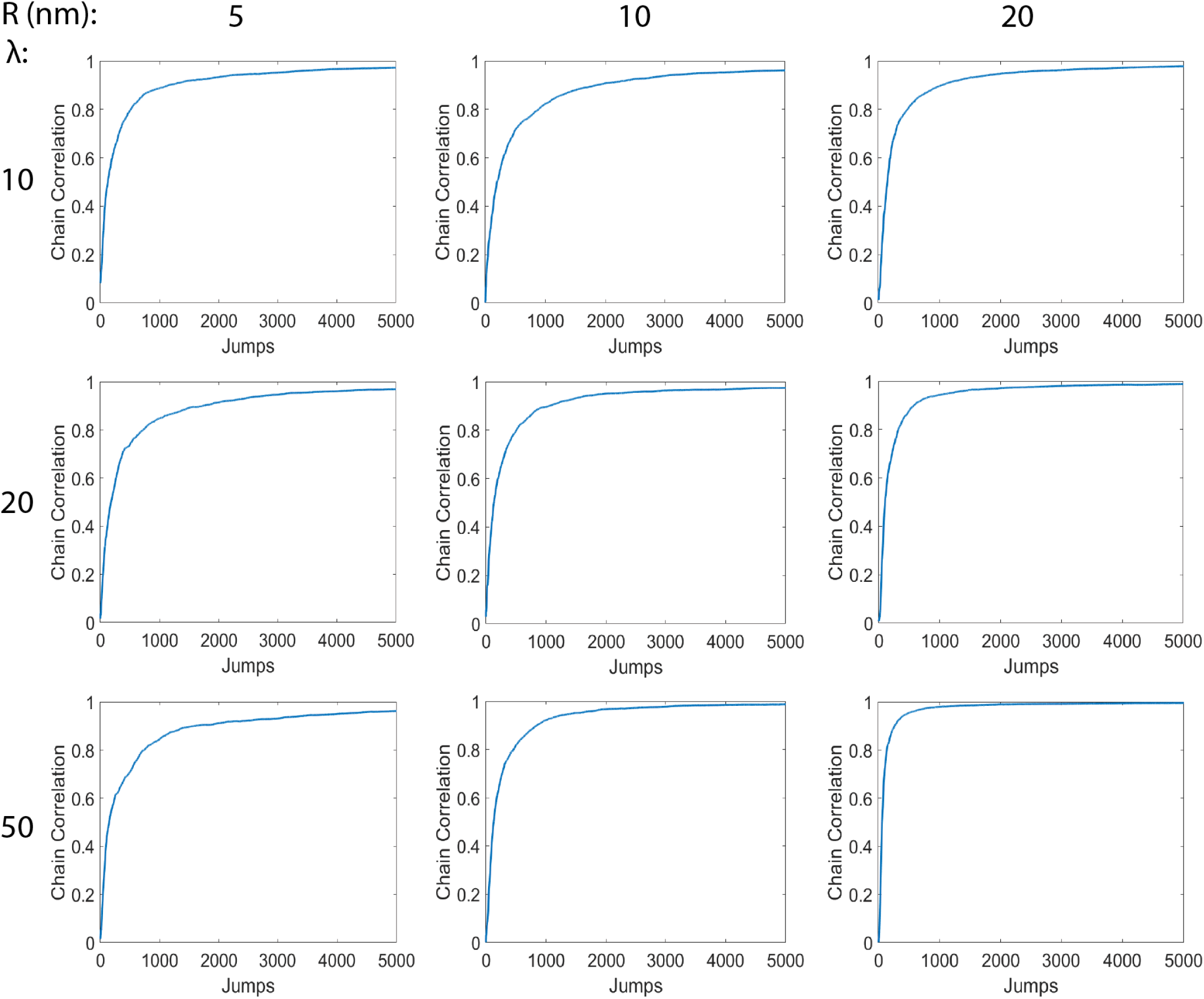
Chain mixing and convergence. The posterior image correlation of the chains over time from independent runs averaged over 10 identical 8-mers similar to Supplementary Fig. 3.

**Supplementary Figure 8:**
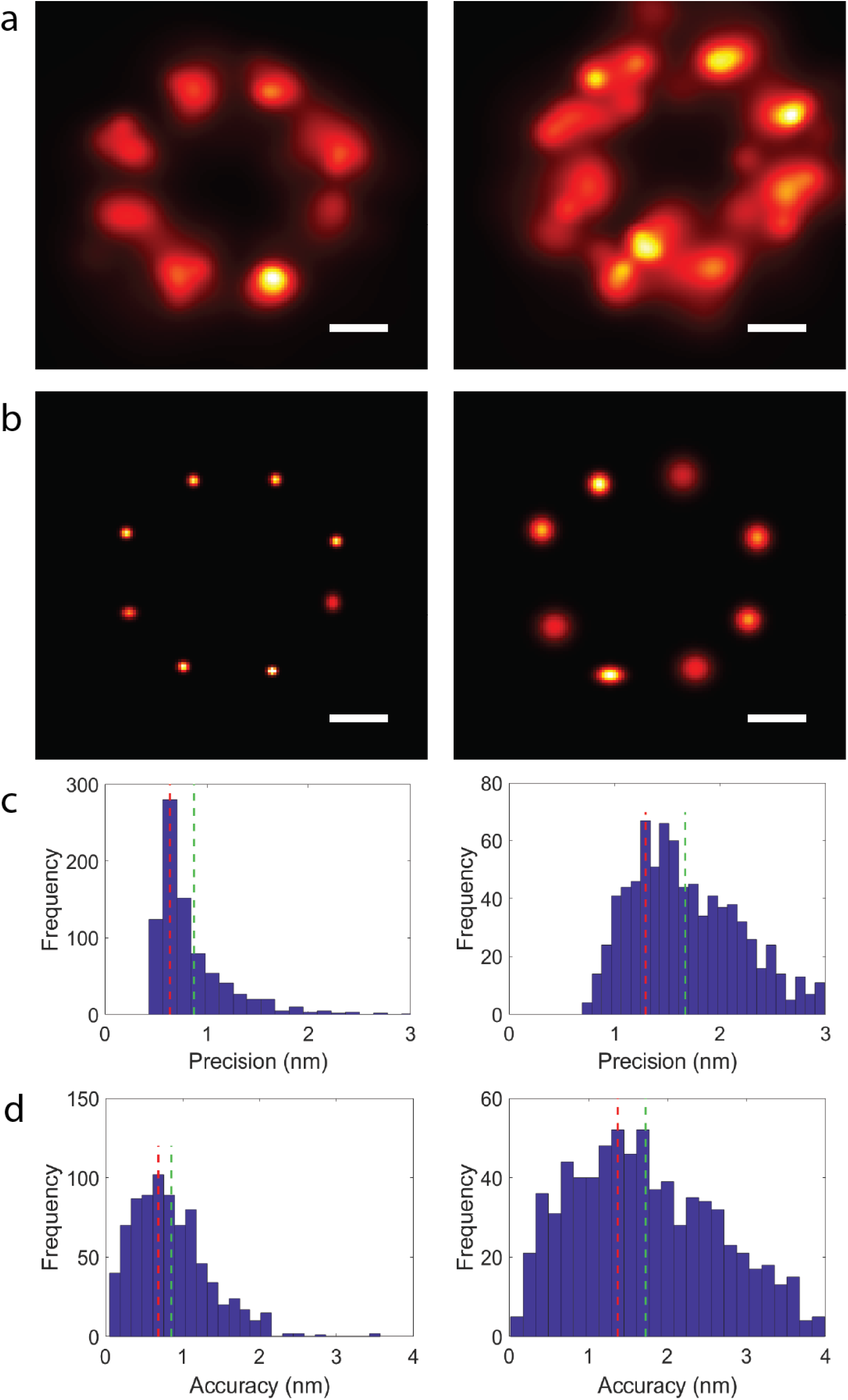
Impact of emitter movements on precision. The emitters in the left column had no movement, while the right column emitters had 10 nm movements over the entire 10,000 frames. The average intensity of binding/blinking events, PSF size, radius of the 8-mers and the average number of blinking/binding events per emitters were 1800 photons, 120 nm, 20 nm and 50, respectively. Row a, b, c and d are the super-resolution images, MAPN images, histograms of resulting precisions and accuracies, respectively.

**Supplementary Figure 9:**
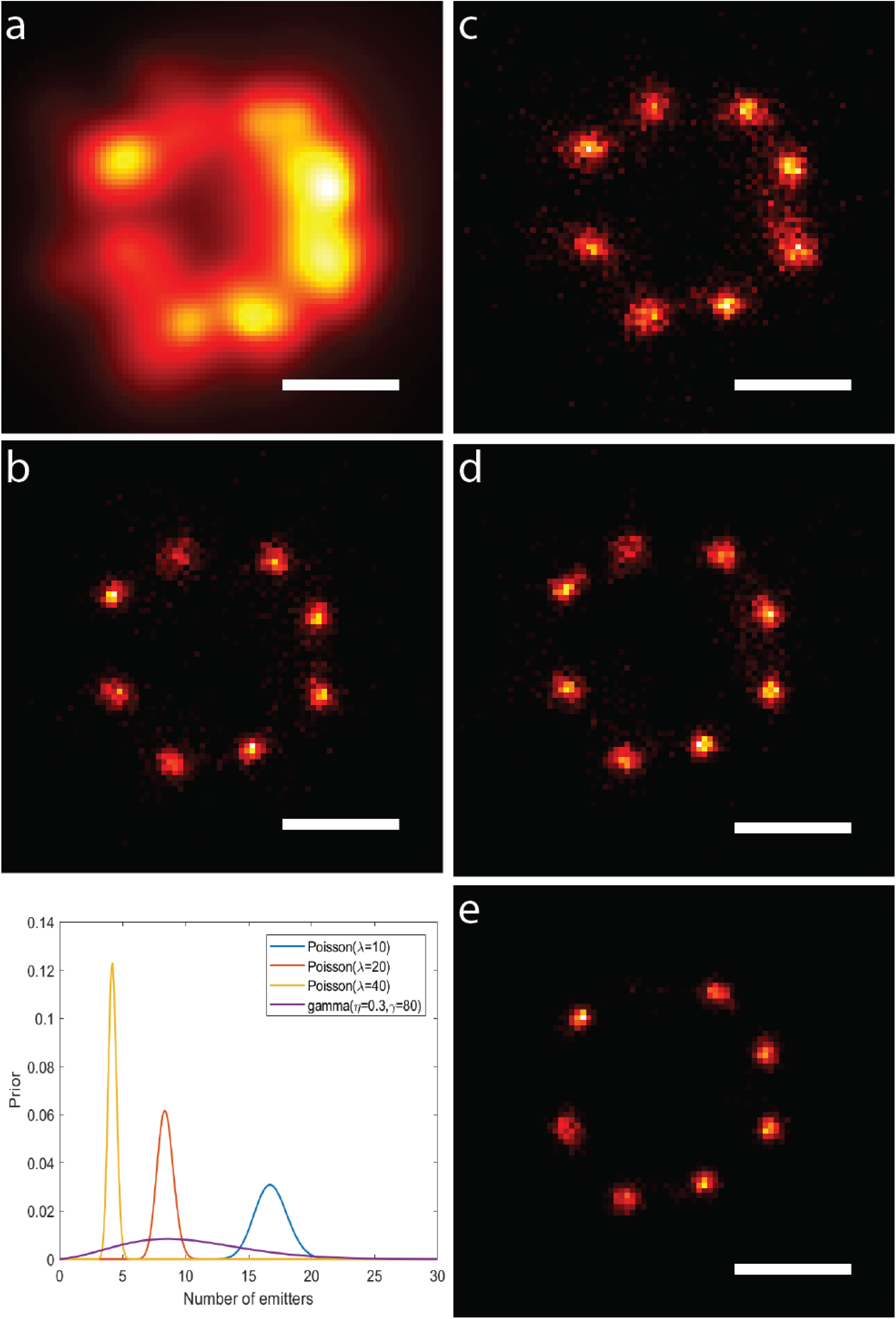
BaGoL performance with different priors on number of emitters. (a) Super-resolution image of an 8-mer, (b) resulting posterior image using a gamma prior with a broad range, *η* = 0.3, *γ* = 80, (c) posterior image using a Poisson prior with *λ* = 10, (d) posterior image using a Poisson prior with *λ* = 20, (e) posterior image using a Poisson prior with *λ* = 40. The plot depicts the four different priors used to process the data. The priors were obtained by considering different average numbers of localizations per emitter, *λ*, where the total number of localizations was fixed. Super-resolution data was simulated with 8 emitters evenly spaced on a ring with *R* = 10 nm and *λ* = 20. The average intensity of each blinking event was 1800 photons with a PSF size of 120 nm. Scale bars are 20 nm.

**Supplementary Figure 10:**
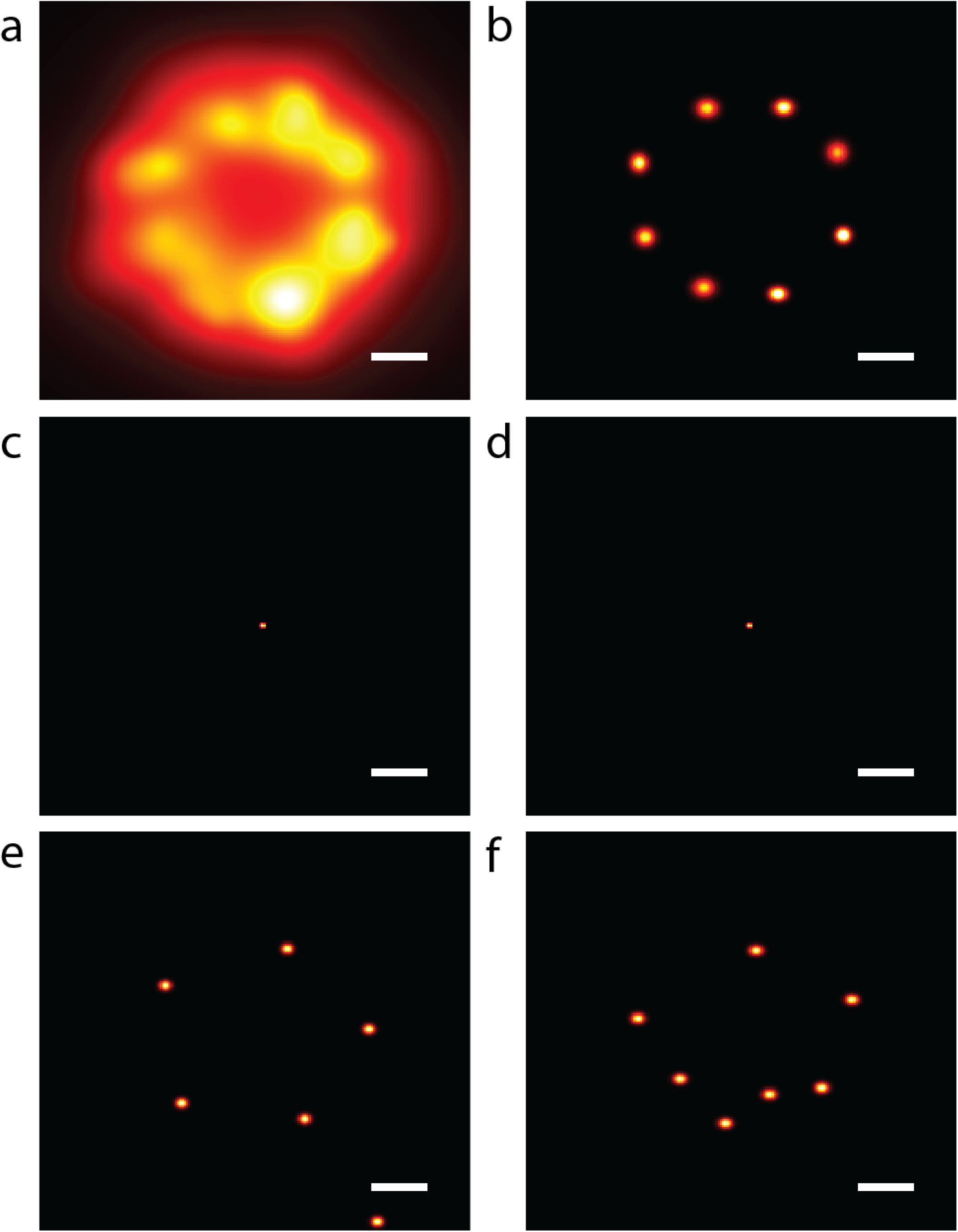
Comparison of BaGoL to other clustering algorithms. Eight emitters were evenly spaced on a ring with radius of 10 nm, an expected *λ* = 50 blinking/binding events per emitter, and an average of 1700 photons per event. (a) Synthetic super-resolution image. (b) MAPN result from BaGoL. (c) Result from the DBSCAN algorithm with the maximum distance between the points within a cluster set to the mean localization precision and 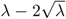 as the minimum number of points within a cluster. (d) Outcome of the algorithm described in [19], where we used the recommended value, 20, for the Dirichlet prior and a gamma prior on the size of the clusters with average of 4 nm, which was the mean value of the localization precisions. (e) The best outcome of *k*-means from 10 different initializations with *K* = 8. (f) Gaussian Mixture Model result using 8 clusters.

**Supplementary Figure 11:**
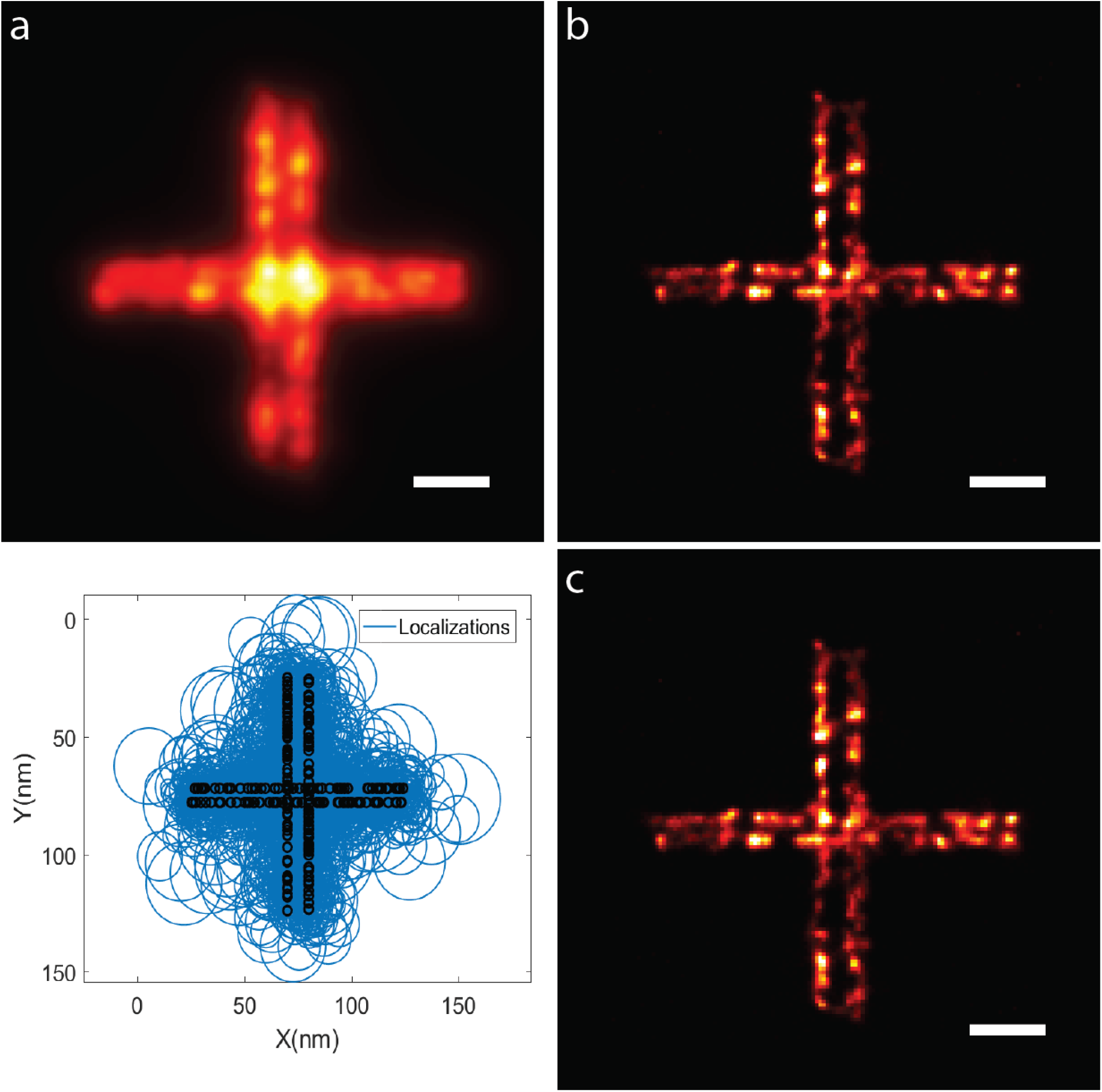
Analysis of continuous lines of emitters. (a) Super-resolution image, (b) posterior image obtained using a Poisson prior for number of emitters, (c) posterior image obtained using a broad gamma prior for number of emitters. In the plot, the blue circles show localizations with radii equal to the precisions, while the black circles show true emitter locations. To simulate this data, 70 emitters were randomly placed on each line of length 100 nm. The horizontal and vertical lines are separated by 6 nm and 12 nm, respectively. The average intensity, PSF size and mean number of localizations per emitter were, respectively, 1800, 120 nm and 20.

**Supplementary Figure 12:**
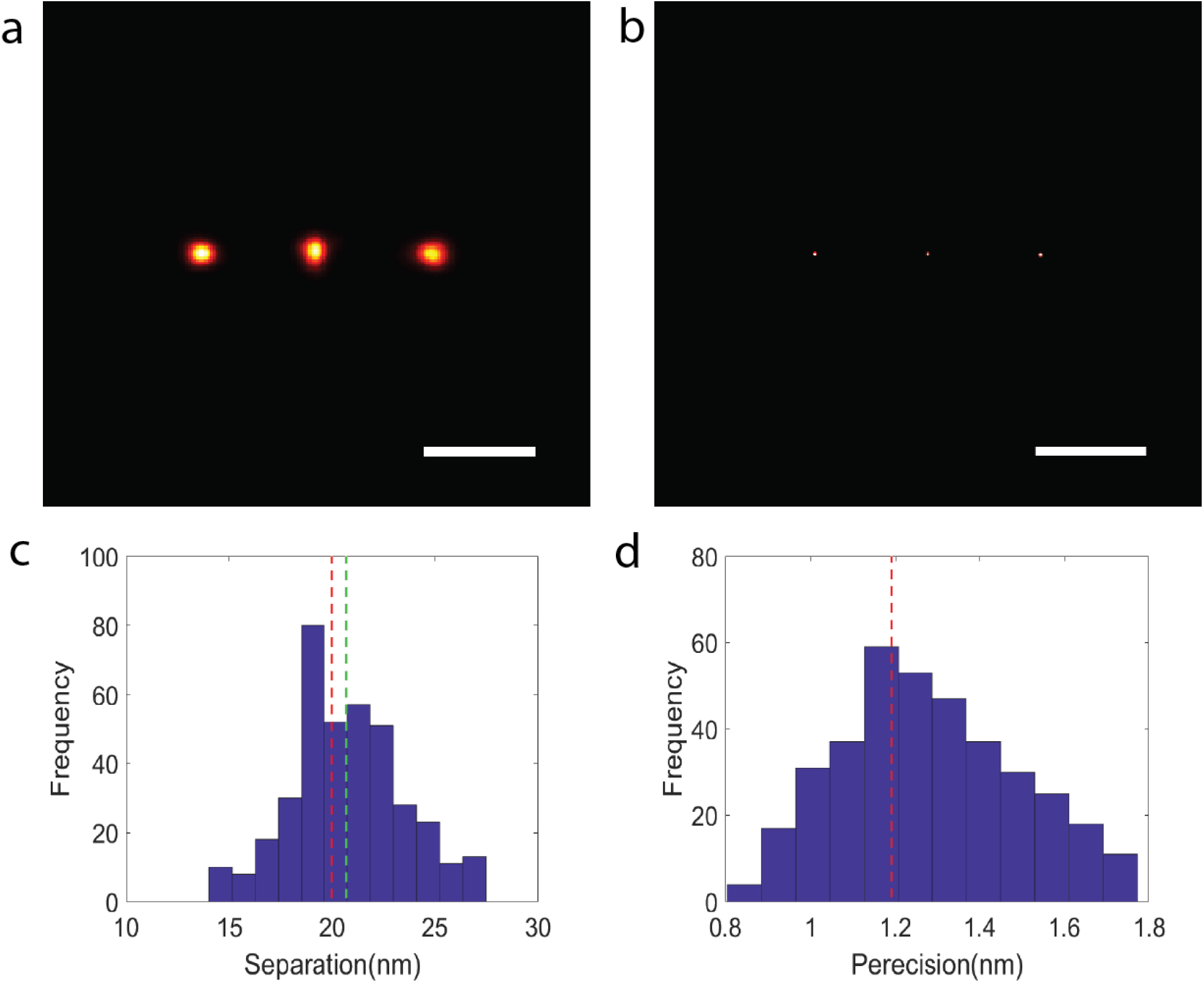
The separation and precision for Gattaquant DNA-ruler. (a) Collection of aligned MAPN results from 150 DNA-rulers. (b) The MAPN result applying BaGoL to the collection of aligned MAPN coordinates in (a). Scale bars are 20 nm. (c) The histogram of separations between the emitters from 150 DNA-rulers. The red line shows 20 nm separation and the green line represents the found separation from (b), 20.7 nm. (d) The precisions for the MAPN localizations from 150 DNA-rulers.

## 1 Supplementary Note 1: Improved Precision from the Combination of Localizations

The data in this problem is a set of *N* localizations, 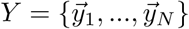 and the uncertainties/precisions attributed to those localizations, 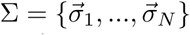. It is assumed that these localizations are from different blinking/binding events, recorded at *T* = {*t*_1_, *…, t*_*N*_}, generated from an underlying set of emitters. The localization precision for the *i*th blinking/binding event in practice is a function of the background and number of photons collected, and would be returned by the fitting algorithm. For the sake of illustration, we can approximate this as [8]

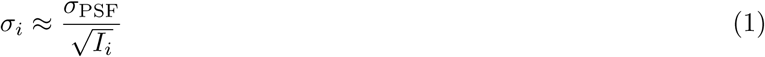

where *σ*_PSF_ and *I*_*i*_ are the size of the point spread function (PSF) and the number of photons from the *i*th blinking/binding event. The goal is to detect and localize those emitters by appropriate grouping and combining of the given localizations. In the grouping stage, an index parameter *Z* is introduced where *Z*_*i*_ = *j* indicates that the *i*th localization is associated to the *j*th emitter. The likelihood of the *i*th localization belonging to the *j*th emitter with location 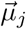 for the *d*th spatial component is given by

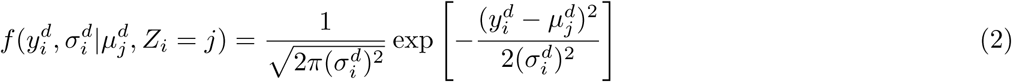

where the upper indices, *d*, indicate the components of 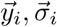 and 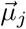. The likelihood of the *j*th emitter given the set of localizations associated to this emitter is

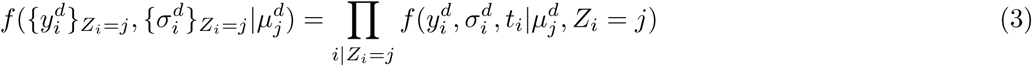

where 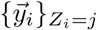 and *i*|*Z*_*i*_ = *j* represent the set of localizations that are associated with the *j*th emitter, and 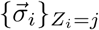 are the uncertainties associated with those localizations. The likelihood (3) can be used to calculate the precision 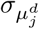 for the emitter location 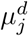 obtained via the combination of the localizations associated with this emitter:

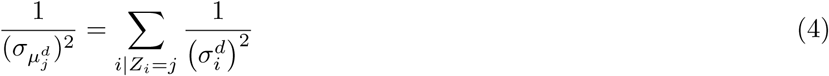

Substituting (1) into (4) yields

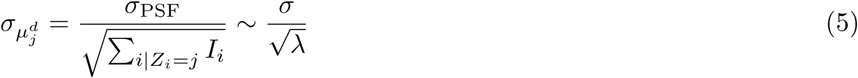

The denominator in (5) is the square root of the sum of all the photons emitted in every blinking/binding event associated to the *j*th emitter during data acquisition. *σ* and *λ*, respectively, indicate the average localization precision and the average number of blinking/binding events per emitter. Therefore, the combination of the blinking/binding events results in a better localization precision, roughly by a factor of 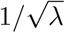.

We allow independent movement for each emitter over time to accommodate factors such as residual drift, alteration in polarization direction, bad fixation, etc. Including these movements as a linear drift, (2) takes the for

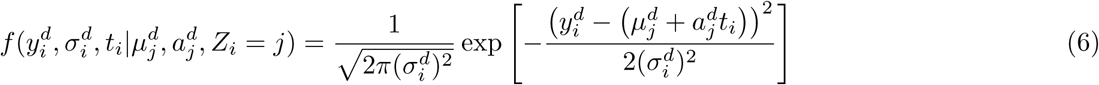

in which 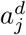 and *t*_*i*_ are the *d*th component of the movement per frame (velocity) of the *j*th emitter and the occurrence time (frame number) of the *i*th blinking/binding event. Taking into account these movements, the likelihood for the *j*th emitter takes the following form

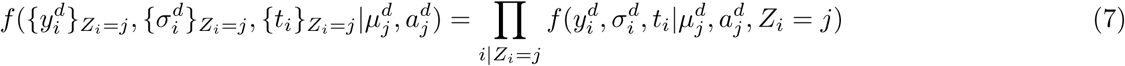

The likelihood (7) can be employed to calculate the values and precisions for *µ*_*j*_ and *a*_*j*_ in the presence of the movements. The observed information is given by

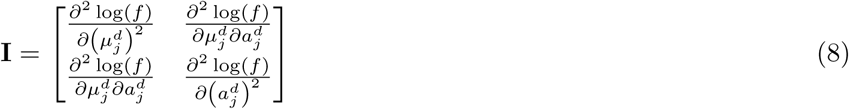

where *f* is the likelihood given in (7). Let **Ξ** be the covariance matrix given as the the inverse of the information:

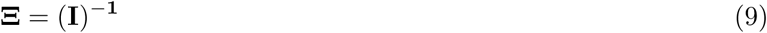

The maximum likelihood estimates are

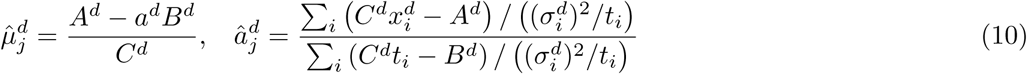

The square root of the diagonal elements of **Ξ** give the precisions

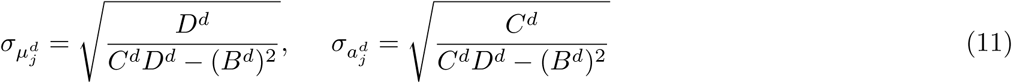

where

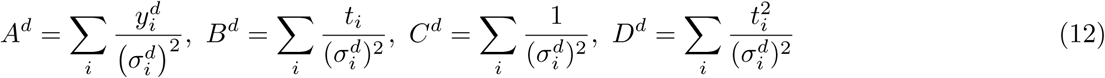

and the sum is over the localizations associated to the *j*th emitter. Note that the presence of emitter movements has a negative impact on the localization precision. This can be shown by comparing the moved emitter precision (11) with the static emitter precision (4)

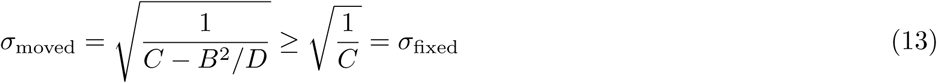

where the super-scripts are dropped. Note that *B, C* and *D* are positive parameters.

## 2 Supplementary Note 2: Reversible Jump Markov Chain Monte Carlo

Reversible Jump Markov Chain Monte Carlo (RJMCMC) [11, 12, 27] is an extension of Markov Chain Monte Carlo (MCMC) [13, 14] that allows sampling from parameter spaces with varying numbers of parameters, thus making inferences about the number of parameters as well as the parameters themselves. For the grouping problem that we will address here, the number of parameters is related to the number of emitters, *K*, the parameter 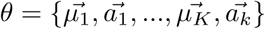 corresponding to the true emitter positions and their corresponding movements, and the latent variable *Z*, which represents assignments of localizations to different emitters. There are 4 different types of jumps in BaGoL (Fig. 1). ***Add*** explores the possibility of adding a new emitter to the current model. ***Remove*** is the opposite of add and examines the feasibility of eliminating one of the existing emitters from the current model. ***Allocate*** uses the Gibbs algorithm [27, 28] to propose an allocation parameter *Z*. ***Move*** updates the locations and movement parameters, *θ*, employing the Gibbs procedure.

### 2.1 Likelihood, Prior and Posterior

RJMCMC is often used in a Bayesian context to sample from the joint posterior of a system *π*(*K, θ*_*K*_|*Y*), where both the number of parameters and the parameters themselves are unknown.

#### 2.1.1 Posterior

In our problem, we take the posterior, *π*(*K*, (*θ*_*K*_, *Z*)|*Y*, Σ, *T*), which is proportional to the product of the likelihood, *P* (*Y*, Σ, *T* |*K, Z, θ*), and priors, *P* (*K*), *P* (*Z*|*K*) and *P* (*θ*_*K*_|*K, Z*)

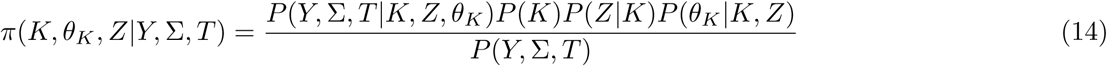

where 1*/P* (*Y*, Σ, *T*) is the constant of proportionality which is called the evidence.

#### 2.1.2 Likelihood

The likelihood of the data can be written as

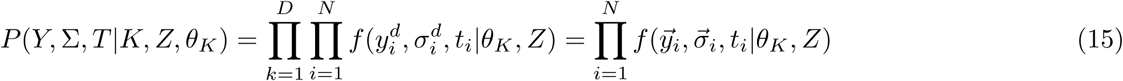

where *D* is number of the spatial components and 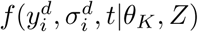 is given in (2, 6) for the static and non-static emitters as appropriate.

#### 2.1.3 Prior Distributions

The prior on the number of emitters is calculated assuming a Poisson or gamma distribution of localizations from each emitter with a mean number of localizations per emitter, *λ*. Given a total number of observed localizations *N*, the prior is then

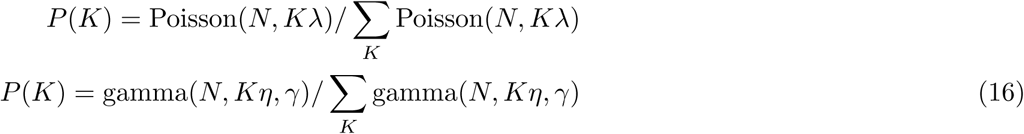

where *γ* is the scale parameter and *ηγ* = *λ*. The prior on allocations is taken to be a multinomial distribution representing *N* localizations distributed among *K* emitters, with the same probabilities, 1*/K*, where |{*Z* = *j*}| is the number of localizations allocated to the *j*th emitter.

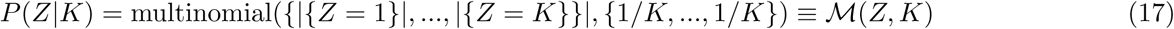

The prior on emitter positions is taken as a uniform distribution over the length of the range of the data in each dimension, *R*^*d*^.

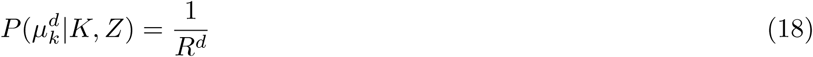

The prior on the drift parameters is taken as a uniform distribution over a maximum allowed drift, *a*_0_.

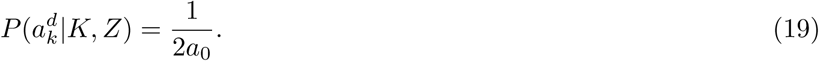

### 2.2 Jump Types

There are two main types of jumps used by BaGoL: two intra-model jumps called ***Allocate*** and ***Move***, and two complimentary inter-model jumps called ***Add*** and ***Remove***. The intra-model jumps allow the exploration of the parameters of the current model, which are the emitter locations, drift velocities, and assignment of the localizations to the current emitters. The inter-model jumps let the chain move between models with different numbers of emitters.

#### 2.2.1 Proposing a Jump

At each step in the chain, we pick a random jump type using an occurrence probability of each jump given by the user, which we call *P*_All_, *P*_M_, *P*_Add_ and *P*_R_, respectively, for the probabilities of proposing an ***Allocate***, a ***Move***, an ***Add*** and a ***Remove*** (for instance, *P*_All_ = 0.3, *P*_M_ = 0.3, *P*_Add_ = 0.2, *P*_R_ = 0.2). Note that we have

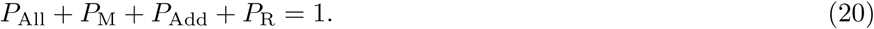

Each jump is accepted or rejected as specified for each jump below.

#### 2.2.2 Intra-model jumps

##### Move

The new values for emitter locations and their drift velocities (if used) are updated using a Gibbs update. The new parameters are taken from the conditional distribution, which is proportional to the likelihood and has the form of a multivariate normal distribution

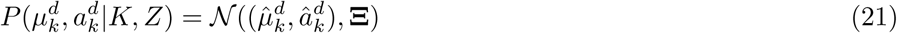

where 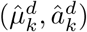 are the maximum likelihood estimates given in (10), and **Ξ** is the covariance matrix given in (9). If the proposed move is outside the range allowed by the uniform priors on *µ*^*d*^ and *a*^*d*^, the jump is rejected, otherwise it is accepted. During a ***Move***, all 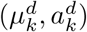 pairs are updated independently using fixed *K, Z*.

##### Allocate

Assignments of localizations to emitters are updated by sampling from

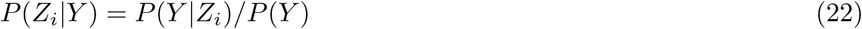

and accepting the complete set of test allocations *Z*′ with probability

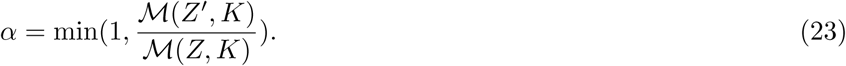

where ℳ represents the multinomial distribution.

#### 2.2.3 Inter-model jumps

***Add*** and ***Remove*** are complementary reversible processes. They allow the addition and removal of an emitter. The proposed jumps are accepted with the probability

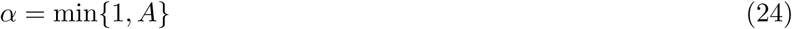

where

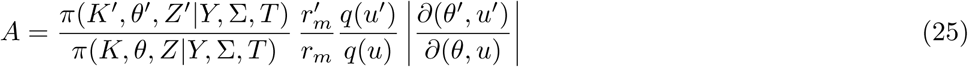

and the prime indicates the proposed parameters. The first term is the ratio of the posteriors. 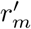 and *r*_*m*_ are the probabilities of proposing a jump and its complementary jump. *q*(*u*) is the proposal distribution corresponding to the probability of proposing model *θ*′ given *θ* with *q*(*u*) being the inverse. The parameter *u* is an auxiliary parameter used together with *θ* to generate *θ*′. The last term is the Jacobian of the transformation from (*θ, u*) to *θ*′.

##### Add

The position and drift velocity of a new emitter is selected by drawing a sample from the priors on these parameters. Allocations are made using the new set of emitter locations and drift velocities via (22). The jump is accepted with probability given by (24), where

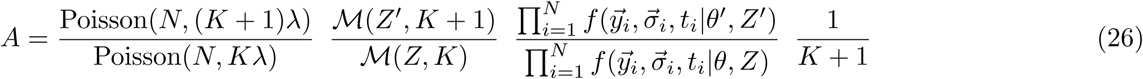

where the ratios from left to right are the prior ratio on the number of emitters, the prior ratio on the allocation distribution, the likelihood ratio, and a ratio that accounts for the probability of this emitter to be chosen for deletion on the reverse jump ***Remove***. Because the proposal distribution *q*(*µ*) for the new parameters is taken from the prior distributions on *θ*, this factor and the prior ratio on *θ* cancel and do not appear in (26). Since the current emitters and drift velocities, *θ*, are held fixed and *u* is itself the location and drift for the new emitter, the Jacobian is equal to one.

##### Remove

One of the existing emitters is selected at random and deleted from the model and the localizations are allocated across the remaining emitters as described in the ***Add*** section above. The acceptance probability is given by

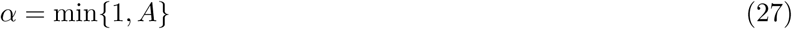

where the ratio A is given by

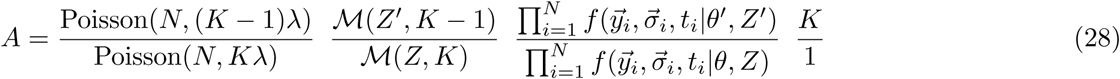

## 3 Supplementary Note 3: Chain Setup and Generation

The chain is initialized to a state with the number of emitters ∼ *N/λ* and random emitter positions selected from the position prior. The jumps described in Supplementary Note 2 are iteratively proposed within a loop. If jumps are accepted, *Z*′, *θ*′ (the proposed values) are recorded in the chain, otherwise *Z, θ* (the current values) are recorded. The chain is comprised of two parts. The beginning portion of the chain is called the burn-in when the chain has not yet settled to a stationary distribution. In the second portion, the chain is exploring the stationary posterior distribution; this portion of the chain is returned for further analysis.

## 4 Supplementary Note 4: Data Flow

Before processing the data using BaGoL, some pre-processing is required. The localizations are frame connected across consecutive frames to connect the localizations that are attributed to a single blinking/binding event that spans more than one frame. Next, the blinking/binding events that are too bright are considered outliers [16] and removed from the inputs to BaGoL. The BaGoL algorithm has several stages. Data often is generated over a large field of view (e.g., a whole cell), whereas this method is designed to be applied to small clusters of emitters. First, the region of measurement containing all the localizations is split into smaller sub-regions. Neighboring sub-regions are overlapped to avoid edge artifacts. Second, the localizations that are considered spurious are filtered out using a nearest neighbor algorithm. A localization is considered signal when it has at least *N* neighbors within *R* distance. Parameters *R* and *N* are specified by the user and can be estimated by inspecting nearest neighbor distributions in the data. The localizations in each sub-region are then further separated into discrete clusters using the hierarchical clustering algorithm [20]. This algorithm is very fast, and has only one parameter (the maximum distance between localizations within a cluster). Each cluster is then processed by the core RJMCMC algorithm. The post-burn-in portion of the chain is returned for further analysis. The chains inside the overlapping areas are eliminated and the chains from the pre-clusters are stitched back together to obtain the posterior for the entire data. In addition, we extract the states with the most repeated model (MAPN) from the returned chain and use it to calculate the MAPN parameters. The extracted states have the same number of emitters, but the emitters are ordered differently and their positions are also slightly altered in each state. Therefore, we employed *k*-means clustering to find the emitter locations and the number of localizations per group. After the localizations are found, then the emitters within overlapping regions are removed to avoid counting an emitter twice, and the list of MAPN coordinates and precisions are returned.

